# PGE2 modulation of stimulus valence directs zebrafish behavioral fever

**DOI:** 10.64898/2026.05.08.723740

**Authors:** Bradley Cutler, Claire E. Tobin, Hailey L. Hollins, Kaarthik A. Balakrishnan, James A. Gagnon, Martin Haesemeyer

## Abstract

Animals seek warmth during fever, but whether this behavior reflects altered temperature perception or interpretation remains unresolved. A prevailing view holds that fever reduces perceived temperature, making the environment feel colder. Here, we use behavior, modeling, and calcium imaging in larval zebrafish to test these hypotheses. We find that viral immune signals drive larval zebrafish to seek warmer temperatures in a PGE2-dependent manner. We localized PGE2 receptors to thermoregulatory brain regions that are conserved across vertebrates. Through comparative behavioral modeling we show that contrary to the prevailing view, fever does not reduce the perception of warmth. Instead, PGE2 alters the valence of temperature changes, such that heating becomes attractive while cooling turns aversive. Calcium imaging reveals that modulation of medullary activity by immune signals directs these changes in stimulus valence. Together, these results identify PGE2-driven changes in stimulus valence as a key principle of the sensorimotor basis of fever.

## Introduction

During physiological fever, organisms raise their body temperature to assist in fighting an infection ^1–7^. Fever is conserved in most endothermic and ectothermic animals from arthropods to humans ^5,8^. Endotherms can metabolically raise their body temperature under autonomous control ^9^. However, autonomous thermoregulation is energetically costly ^10^. Therefore, even endotherms will use behavioral means to elevate their body temperature to minimize autonomous regulation ^11–14^. This phenomenon is referred to as behavioral fever, and it is the dominant mechanism of fever induction in ectotherms ^8^.

The molecular cascades involved in the generation of fever have been well studied, especially in mammals. As a final step, prostaglandin E2 (PGE2) signals through EP2 and EP3 receptors to induce fever ^15–23^. Command neurons inducing sickness behaviors that express EP2 or EP3 receptors have been located within the mouse preoptic area (POA) ^14^ and nucleus of the solitary tract ^24^. However, we fundamentally do not understand how vertebrates seek warmer temperatures during fever. A prevailing view is that immune signals reduce the perception of warmth ^9,25^. This illusion of cold then drives animals to seek warmer temperatures. Alternatively, immune signals might change the interpretation of temperature stimuli and thereby increase cold aversion without altering the perceived temperature. These competing hypotheses have not been tested, limiting our understanding of the influence of immune signals on behavioral thermoregulation. This is partially because we lack understanding of the circuits that control the seeking of preferred temperatures and, to a larger extent, because it is unclear how circuits and behavior are modulated by fever signals. To gain such understanding requires observation of neural activity in intact animals, which is difficult in mammalian models due to the limited access to subcortical structures. Here, we established a behavioral fever paradigm in larval zebrafish to overcome these limitations. Larval zebrafish are optically transparent which allows us to perform calcium imaging at cellular resolution across entire brain regions. At the same time, prior work identified the behavioral and neural programs zebrafish larvae use to thermoregulate ^26–28^.

Here, we show that larval zebrafish will seek out temperatures that are about 1 °C warmer than their baseline preference after a viral immune challenge. This increase in preferred temperature is comparable to behavioral fever in mammals ^14,29^ and, as in mammals, it critically depends on PGE2 signaling. In line with this finding, analysis of scRNAseq data localizes canonical immune signaling cascades to hypothalamic endothelial cells. We furthermore identify expression of PGE2 receptors in conserved thermoregulatory brain regions. This establishes behavioral fever in larval zebrafish as a general vertebrate system to study the basis of warmth seeking during an immune challenge. Comparison of swim kinematics in the baseline and fever state argues against the prevailing hypothesis that fever is triggered by a reduction in the perception of warmth. Probabilistic models of thermoregulation instead demonstrate that immune signals alter the valence of temperature change stimuli. Namely, viral pathogen associated molecular patterns (PAMPs) make heating stimuli more attractive while cooling stimuli become aversive. Functional calcium imaging in the medulla of larval zebrafish identifies changes in the activity of neurons encoding heating and cooling stimuli that mirror the observed changes in valence. Taken together, these findings support a model in which distributed PGE2 signaling modulates medullary temperature encoding to alter the valence of temperature stimuli leading to larval zebrafish seeking out warmer temperatures (behavioral fever). We hypothesize that this change in stimulus valence, rather than a reduction in perceived warmth, is a conserved feature of behavioral fever.

## Results

### Viral PAMPs induce behavioral fever in larval zebrafish

We set out with the goal of using larval zebrafish to understand computational principles of behavioral fever in vertebrates. While fever has been described in adult fish ^4,6,31,32^, the ability of larval zebrafish to develop behavioral fever has been contested ^33^. Therefore, we first assessed whether larval zebrafish exhibit phenotypical behavioral fever when PAMPs are detected by the immune system. To this end, we developed an assay using the dsRNA, high molecular weight poly(I:C), as a viral PAMP. We injected poly(I:C) together with a fluorescent tracer in PBS into the tail vein of larval zebrafish to induce systemic inflammation. After letting the animals rest for 16 hours, we placed them into a custom-designed thermal chamber which allows us to establish linear temperature gradients using Peltier cooling and heating ^27^. Specifically, we generated a horizontal temperature gradient from 20 °C on one end of the chamber to 30 °C on the other (Figure 1A). We tracked the heading and position of each larva at 100 Hz to capture detailed behavioral kinematics while it explored the gradient for 30 minutes (Figure 1A).

**Figure 1:**
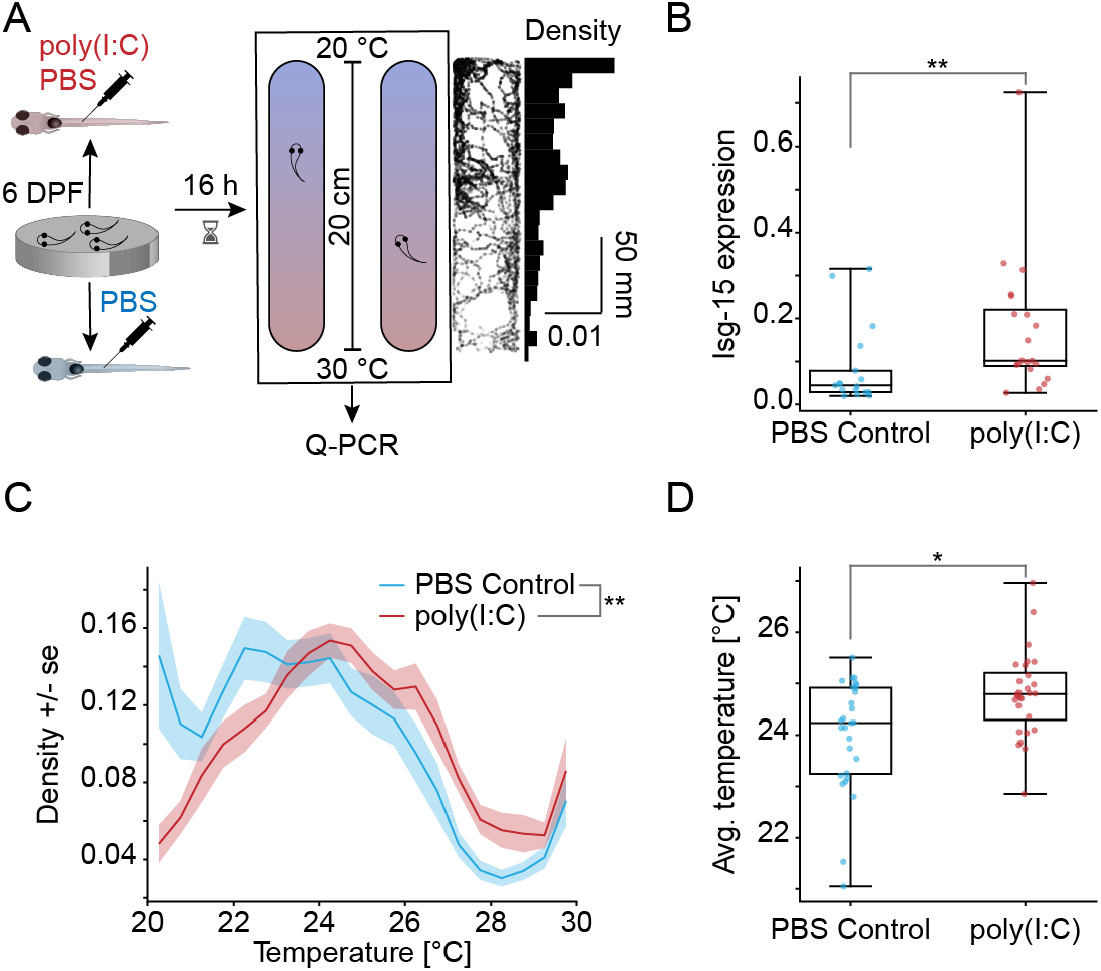
Behavioral Fever in larval zebrafish. **A**: Schematic of the assay and experimental timeline. Plots on the right depict the trajectory of one example control fish across an entire experiment as well as its occupational density within the temperature gradient. **B**: Isg-15 expression level as a fraction of Mob4 expression across treatments. ∗∗ : *p <* 0.01 Mann-Whitney-U test across larvae. **C**: Densities of time spent at different temperatures in the gradient for PBS-injected controls (blue) and poly(I:C)-injected fish (red). Shading indicates bootstrap standard error across larvae. ∗∗ : *p <* 0.01 bootstrapped KS-test ^30^ across larvae. **D**: Average experienced temperature in the gradient across experimental time for each treatment. ∗ : *p <* 0.05 Mann-Whitney-U test across larvae. See Table S1 for counts and exact p-values. Also see Figure S1

We and others recently demonstrated that viral infections induce robust immune responses in larval zebrafish, including expression of the interferon response gene *isg15* ^34,35^. To confirm that poly(I:C) injection is sufficient to drive an immune response, we used RT-qPCR to measure relative *isg15* mRNA abundance in individual larvae after the behavioral assay (∼ 24 hours after injecting poly(I:C)). We found that poly(I:C) injected fish showed a significant increase in relative *isg15* mRNA abundance when compared with controls injected with only PBS (Figure 1B). This indicated that larval zebrafish exhibited a viral inflammatory response after injection of poly(I:C).

We then tested the hypothesis that larval zebrafish would develop behavioral fever after poly(I:C) injection. Analyzing time spent at different temperatures within the gradient revealed an increase in the preferred temperature. Specifically, after poly(I:C) injection, larval zebrafish spent more time at temperatures above 26 °C and less at temperatures below 23 °C (Figure 1C), leading to a significant shift in the occupancy distribution to warmer temperatures compared to PBS injected controls (Figure S1A). While temperature preferences were variable across individual fish, shifts in occupancy within the gradient could also be observed at the level of individual larvae (Figures S1B-C). Overall, poly(I:C) injection led to a significant increase of almost 1 °C in the average experienced temperature relative to controls (Figure 1D), which is similar to preference shifts observed during behavioral fever in mammals ^14,29^. This demonstrates that larval zebrafish exhibit canonical behavioral fever after injection of a viral PAMP.

Inflammation causes broad sickness behaviors ^36^ and can reduce locomotion ^37,38^. Since our assay relies on larvae navigating towards their preferred temperature, we wanted to ensure that exploration of the chamber was not impaired after poly(I:C) injection. Quantifying per-minute exploratory distances across time and larvae revealed that they do not fatigue over the course of the experiment (Figures S1D-E) and importantly that poly(I:C) injection does not affect exploration within the gradient chamber (Figure S1F). This indicates that while poly(I:C) increases the preferred temperature, it does not impair movement of larval zebrafish on longer timescales. In summary, these data show that poly(I:C) injection drives larval zebrafish to seek out warmer temperatures. This indicates that immune signals in response to viral stimuli alter temperature navigation behavior to favor higher temperatures, i.e. develop fever.

### Zebrafish larvae express genes necessary for immune regulated PGE2 signaling

Work across vertebrates and especially mammals identified key signaling cascades that translate inflammatory immune signals into autonomous and behavioral components of fever (Figure 2A). Binding of cytokines such as interleukin 6 (IL6) or TNF*α* (TNFa) to their cognate receptors on brain endothelial cells leads to the production of PGE2 through enzymatic action of COX-2 and PTGES^40–45^. PGE2 subsequently binds to EP2 and EP3 receptors to induce behavioral and autonomous components of fever ^14,46–49^. EP2 and EP3 receptors are differentiated by their downstream signaling, where EP2 receptors are predominantly thought to excite neurons, while EP3 receptors largely have inhibitory effects with a few exceptions ^50,51^. Since larval zebrafish developed behavioral fever in response to viral antigens (Figure 1) we tested whether elements of this canonical immune signaling cascade are present in the brains of larval zebrafish.

**Figure 2:**
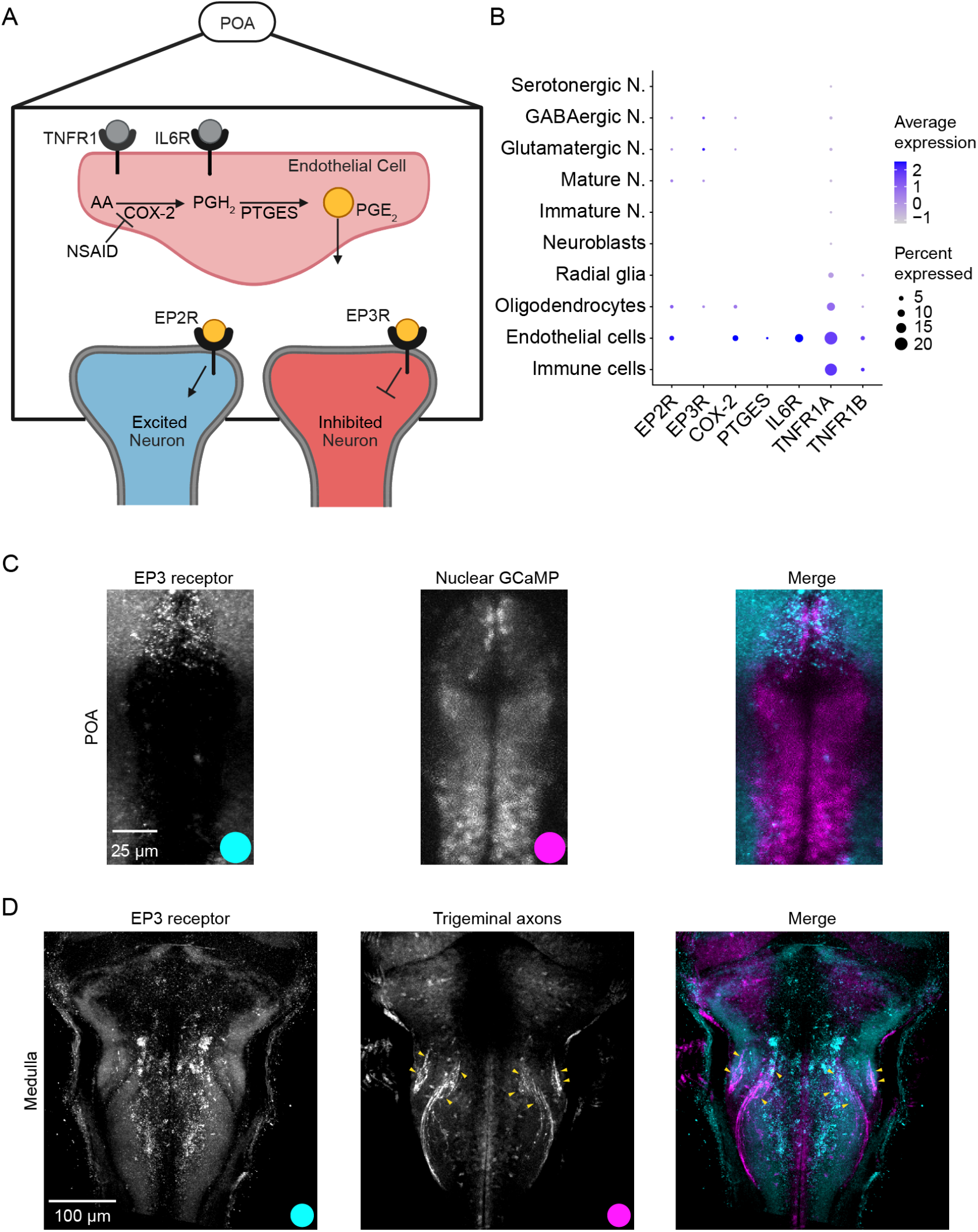
PGE2 signaling cascades are conserved in larval zebrafish. A: Simplified schematic illustrating relevant genes and signaling cascades in mammalian behavioral and autonomous fever. We note that EP3 receptors are predominantly considered inhibitory, however, the gamma isoform can act as both an excitatory and inhibitory receptor via differential G-Protein coupling. B: Dot-plot of immune response relevant genes from re-analysis of Shafer et al. ^39^ dataset. Dot size represents the percentage of neurons expressing the gene while color indicates log of expression level. Dots are thresholded at a minimum of 0.5 % cells that express the gene. C: HCR *in situ* revealing expression of EP3 receptors in POA neurons. Left panel: Signal of EP3 receptor probe averaged across 3 registered brains, middle expression of Elavl3-H2B:GCaMP6s marking neuronal nuclei, right panel is merge of both channels. Colored dots indicate the color assigned to each signal in the merge. D: HCR *in situ* revealing expression of EP3 receptors in the medulla. Left panel: Signal of EP3 receptor probe averaged across 3 registered brains, middle photo-activatable GFP labeling of trigeminal ganglion sensory neurons revealing location of their projections ^27^, right panel is merge of both channels. Colored dots indicate the color assigned to each signal in the merge. Yellow arrowheads indicate terminations of trigeminal fibers in the middle and right panel. See Table S3 for official zebrafish gene names associated with common names used in the figure. Also see Figure S2

We re-analyzed a published scRNAseq dataset that was generated from hypothalamic dissections of 2-week old larval zebrafish brains ^39^. Genes involved in the immune regulated production of PGE2 are expressed in hypothalamic endothelial cells. Focusing on neurons, genes encoding the EP2 and EP3 receptors were expressed in a small fraction of neurons within distinct neural classes (Figure 2B).

To confirm that PGE2 receptors were expressed within the zebrafish POA, as in mice ^14,51^, we used HCR *in situ* staining to visualize EP2 and EP3 receptor gene expression across the zebrafish brain, which we subsequently registered to a common reference (see Methods). We identified strong EP3 receptor expression in the anterior POA of larval zebrafish (Figure 2C), while there was no signal in controls (Figure S2A). The EP2 receptor was also expressed within the POA (Figure S2B), albeit seemingly at a lower level than EP3 receptors. Aside from the expected expression within the POA, we also found strong expression of EP3 receptors within the zebrafish medulla (Figure 2D). There was no signal within the medulla in controls and EP2 receptor expression was absent as well (Figures S2C-D). The medullary EP3 receptor expression was found in neurons near terminal branches of central axons of sensory trigeminal ganglion neurons (Figure 2D middle and right). This suggests a secondary, direct pathway of PGE2 signaling in fever possibly acting in parallel to POA mediated inflammatory signals. Similarly, a more direct action may occur via PGE2 signaling in the right dorsal habenula. The habenula has been implicated in thermoregulation in mice and zebrafish ^28,52^ and we identified expression of EP3 receptors in neurons of the right dorsal habenula (Figure S2G).

Together, these data show that genes involved in canonical immune signaling cascades are expressed in endothelial cells and neurons within the larval zebrafish POA. PGE2 receptors are expressed in key areas of the brain involved in thermoregulatory responses, suggesting a possible role for distributed immune signaling in the generation of behavioral fever. This strongly suggests that larval zebrafish share a conserved immune fever pathway with other vertebrates, including mammals.

### Larval zebrafish behavioral fever requires PGE2 signaling

The conservation of mammalian fever components in larval zebrafish motivated us to determine the necessity of PGE2 in larval behavioral fever. To this end, we augmented our behavioral fever assay using the drug Celebrex (Figure 3A), a selective Cox-2 inhibitor ^53^ that abrogates the production of PGE2 (Figure 2A).

**Figure 3:**
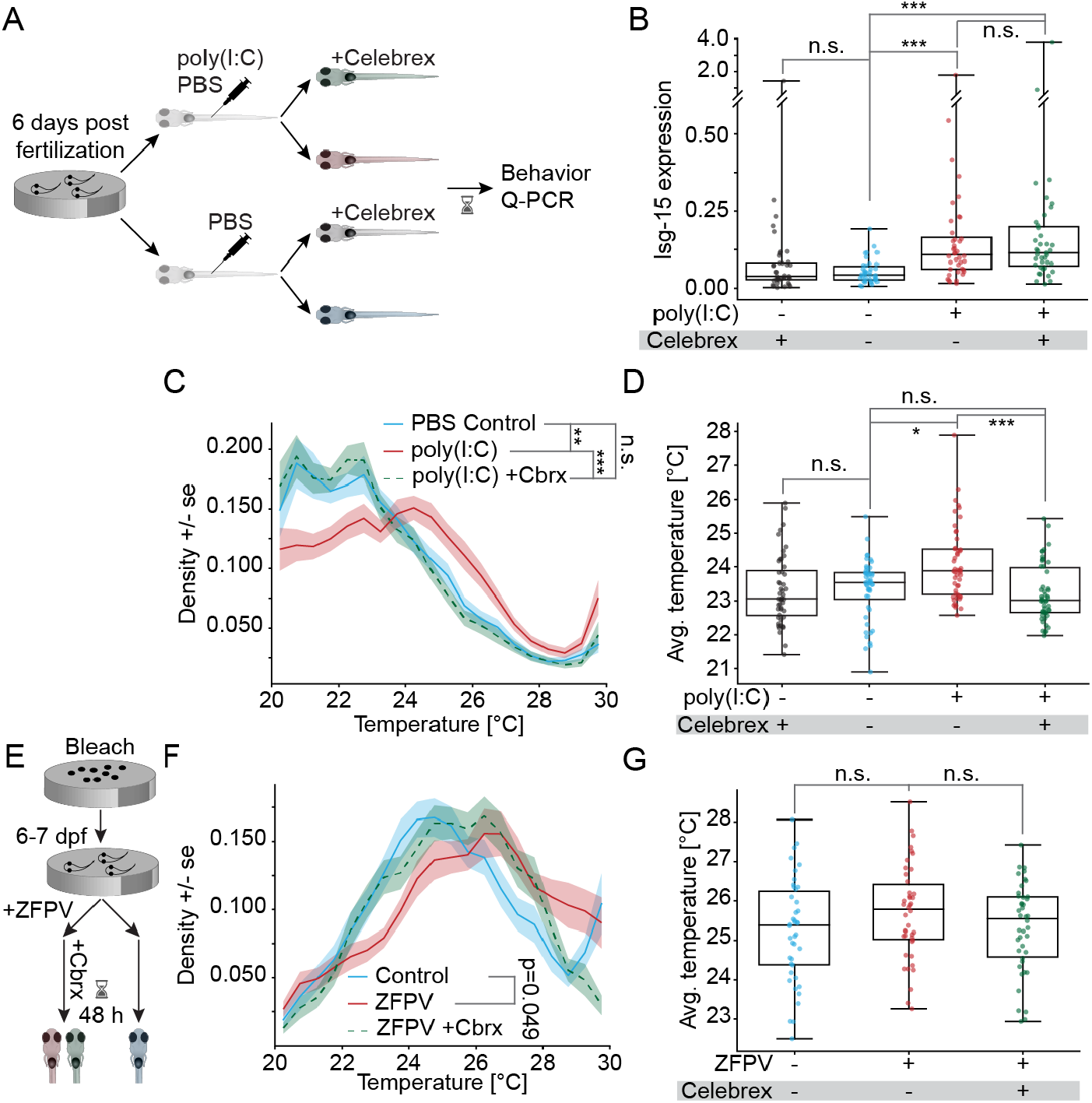
Viral PAMPs trigger behavioral fever in a PGE2 dependent manner. **A**: Schematic of the treatment groups in the Celebrex assay. **B**: Isg-15 expression level as a fraction of Mob4 expression across treatments. Top row below x-axis indicates presence of poly(I:C) in injection, bottom row presence of Celebrex in the bath. Mann-Whitney-U test across larvae. **C**: Densities of time spent at different temperatures in the gradient for PBS-injected DMSO controls (blue), poly(I:C)-injected fish in DMSO (red), and poly(I:C) injected with with Celebrex (green dashed). Shading indicates bootstrap standard error across larvae. Bootstrapped KS-test across larvae. **D**: Average experienced temperature in the gradient across experimental time for each treatment. Mann-Whitney-U test across larvae. **E**: Schematic of the assay and timeline for ZFPV live virus infection experiments. **F**: Densities of time spent at different temperatures in the gradient for Germ-free controls (blue), ZFPV infected fish fish in DMSO (red), and ZFPV infected fish with with Celebrex (green dashed). Shading indicates bootstrap standard error across larvae. Bootstrapped KS-test across larvae. **G**: Average experienced temperature in the gradient across experimental time for each ZFPV treatment. Mann-Whitney-U test across larvae. In all panels, significance after Benjamini-Hochberg correction is indicated as follows. ***: p-value smaller than critical value at 0.1% FDR; **: p-value smaller than critical value at 1% FDR; *: p-value smaller than critical value at 5% FDR; n.s: p-value larger than critical value at 5% FDR. Written out p-values are uncorrected and did not pass criterion after correction. See Table S1 for counts and exact p-values. Also see Figure S3

As before, we injected larval zebrafish with either high molecular weight poly(I:C) dissolved in PBS or PBS alone into their tail vein. After injection, larval zebrafish rested 16 hours in media containing 0.1 % DMSO and Celebrex or only DMSO as a control (Figure 3A). Subsequently, larval zebrafish were assayed in the thermal gradient and RNA was subsequently extracted for RT-qPCR analysis (Figure 3A).

Poly(I:C) led to a significant increase in relative *isg15* mRNA abundance, confirming the induction of viral inflammation in the presence of DMSO (Figure 3B). However, Celebrex did not reduce the interferon response (Figure 3B). This is not unexpected, since the interferon response is triggered by a pathway that is both parallel to the major inflammatory response ^54^ and upstream of the generation of PGE2. This further underlines similarities in zebrafish and mammalian immune signaling pathways.

To test whether PGE2 is necessary for behavioral fever in larval zebrafish, we allowed the animals to freely explore the thermal gradient. Poly(I:C) injection again led to a significant shift in the occupation within the gradient to warmer waters (Figures 3C, S3A-D). This behavioral fever phenotype was reversed by the presence of Celebrex (Figures 3C, S3C-D). The same effect was observed when quantifying the average experienced temperature, which was significantly increased after poly(I:C) injection, an effect that was completely reversed by Celebrex (Figure 3D). In fact, the average temperature after Celebrex administration was slightly lower than baseline conditions. This might suggest a baseline level of inflammation in larval zebrafish, which were not kept in germ free media. Celebrex did not alter gross behavior of larvae as evidenced by the absence of changes to exploratory behavior within the gradient (Figures S3E-H). These results strongly suggest that as in mammals, PGE2 signaling is a necessary step in the induction of fever in larval zebrafish.

As a pure viral PAMP, poly(I:C) elicits a strong and controllable immune response, but is likely simplistic when compared to a true viral immune challenge. To test whether behavioral fever can be induced by a natural viral infection, we turned to the endogenous zebrafish picornavirus, ZFPV^35,55^. ZFPV elicits a strong interferon response, but is non-lethal and only mildly pathogenic ^35^. We treated pathogen-free larval zebrafish with ZFPV for two days before performing our behavioral fever assay (Figure 3E, see Materials and Methods). A control group was exposed to DMSO in addition to a test group being incubated in Celebrex and DMSO for 24 hours. Within the temperature gradient, infection with ZFPV led to an increase in time spent at warmer temperatures, together with a decrease in time spent in cooler areas as expected for behavioral fever (Figure 3F). This was also reflected in a shift in individual fish distributions within the gradient to warmer temperatures (Figures S3I-K). While clearly visible, the overall shift in distribution was not significant after correcting for multiple comparisons (Figure S3L). This is likely a consequence of the known variability of natural infections together with the fact that ZFPV pathology is generally transient and clears on its own ^35^. Application of Celebrex partially rescued ZFPV induced behavioral fever (Figure 3F) bringing the experienced average temperature back to levels of uninfected controls (Figure 3G). As observed with the other treatments, viral infection did not alter larval zebrafish exploratory behavior (Figures S3M-P).

In summary, these data show that behavioral fever in larval zebrafish requires PGE2 signaling both in response to the viral PAMP poly(I:C) and during a natural infection. These data strongly suggest that an immune signaling cascade present in mice ^9,25^ and adult fish ^8^ is active in larval zebrafish.

### Fever does not reduce the perception of warmth

We found that inflammatory stimuli increased the preferred temperature larval zebrafish seek in a PGE2 dependent manner (Figures 1, 3). However, it is unclear how inflammation changes the preference in larval zebrafish or any vertebrate. Broadly, two alternative hypotheses could account for observed changes in preferred temperature. A “perceptual hypothesis” would suggest that the animal perceives temperatures as colder during fever. In this prevalent view, fever arises through a reduction in activity in neurons that signal warmth, which effectively leads to the animal experiencing temperatures as colder than they actually are ^9,14,25^. Under an alternative “setpoint” hypothesis, the perception of temperature would remain unaltered, while the valence of temperature stimuli is adjusted to lead to seeking of warmer temperatures. Both of these strategies would lead to an increase in the preferred temperature the animal seeks. However, the perceptual hypothesis would suggest that all behaviors change as if the temperature is lower (e.g., at 28 °C a larva with fever would behave like a control larva at 27 °C). The setpoint hypothesis on the other hand would allow for selective modulation of behaviors that are important for seeking the preferred temperature. To disambiguate these alternative hypotheses, we analyzed the kinematics of individual swims (Figure 4A) and how they relate to temperature.

**Figure 4:**
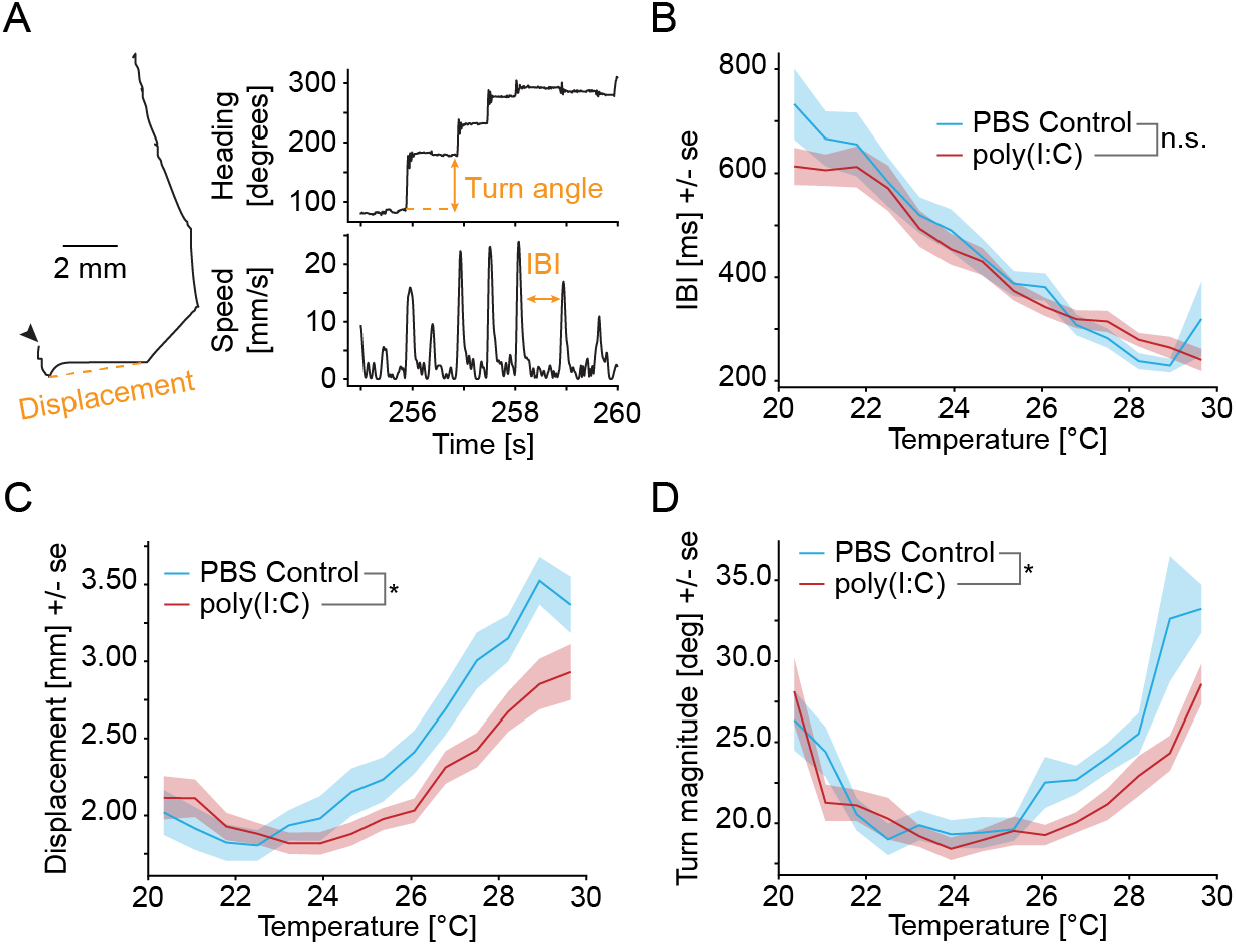
Fever reduces avoidance behaviors. **A**: Example trajectory of a control fish (same as in Figure 1 A) across five seconds (left). Arrowhead indicates start of trajectory. Heading angles (top right) and instantaneous swimming speed (top left) are displayed for the same trajectory. Most important kinematic features analyzed in the following panels are annotated in orange. **B-D**: Average bout kinematics binned by gradient temperature for PBS-injected controls (blue) and poly(I:C)-injected fish (red). Shading indicates bootstrap standard error across larvae. ∗ : *p <* 0.05, *n*.*s*. : *p >* 0.05 bootstrap test of average difference across larvae. **B**: Interbout interval. **C**: Displacement of swim bouts. **D**: Turn magnitude (absolute value of turn angle) of swim bouts. See Table S1 for counts and exact p-values. Also see Figure S4

Larval zebrafish swim in discrete bouts separated by stationary periods, the interbout intervals (IBI) (Figure 4A, bottom right). For each swim bout, we compared the displacement (distance between the start and end point) (Figure 4A, left), the turn angle (Figure 4A, top right), the IBI, and the turn bias (relative turn magnitude after heating versus cooling stimuli, Figure S4E) between poly(I:C) injected fish and PBS injected controls (same data as presented in Figure 1). Within the gradient, both treatment groups experienced the same temperature changes (Figure S4A), in other words, larvae of both groups were subjected to comparable sensory inputs. Overall, the behavioral repertoire of the animal was similar after injection of poly(I:C) or PBS with comparable distributions of swim kinematics (Figures S4 B-D). This is consistent with the lack of change in exploratory movement during fever (Figures S1F and S3H, P).

However, temperature strongly influences swim kinematics in larval zebrafish ^27,56^. We therefore tested whether immune signals change the relationship between temperature and swim kinematics. IBIs decreased almost linearly with temperature and this relationship was unaffected by poly(I:C) (Figure 4B). We specifically note that the shift in the distribution of IBIs in Figure S4B is explained by this lack of change together with the fact that poly(I:C) injected fish spent more time at warmer temperatures. Just like the control of IBI, the control of turn bias by temperature was unaffected during fever. Heating and cooling stimuli bias the magnitude of larval turns and this bias is temperature dependent ^27,28^. Specifically, at cold temperatures, turns after heating stimuli are smaller than turns after cooling stimuli (negative bias). To the contrary, at warm temperatures, turns have a positive bias as they are larger after heating stimuli than after cooling stimuli (Figure S4F). When measuring turn bias in PBS and poly(I:C) injected fish as a function of temperature, there was no difference (Figure S4F). Both the IBI and turn bias results are therefore inconsistent with the perceptual hypothesis, since a reduction in the perception of warmth would be predicted to lead to an overall shift in the relationship between swim kinematics and temperature.

On the other hand, the relationship of temperature and swim displacements was significantly different between conditions (Figure 4C). Specifically, at temperatures above 24 °C the displacement curve for poly(I:C) fish was shifted to the right by close to one degree (Figure 4C). Therefore, poly(I:C) injected larvae at e.g., 28 °C displaced like control larvae at 27 °C. Likewise, poly(I:C) fish showed a significant reduction in the magnitude of turns (absolute turn angle) at warmer temperatures (Figure 4D). This hints at a separation in neural control between swim displacements and turns on the one hand, and IBI and turn biases on the other.

Irrespective of the stimulus modality, increases in displacement and turn magnitude are associated with aversive conditions in larval zebrafish and indicate avoidance behaviors ^27,28,56,57^. These results therefore suggest that larval zebrafish perceived warm water as less aversive after poly(I:C) injection congruent with the fever phenotype. In our Celebrex dataset we observed similar changes in behavior (Figure S4G-J), however there was no significant difference in the modulation of turn magnitude between conditions (Figure S4J). Celebrex partially reversed the effect of poly(I:C) on swim displacements, consistent with the necessity of PGE2 signaling for the fever phenotype.

In summary, these results strongly suggest that inflammatory stimuli do not alter the sensory perception of temperature (perceptual hypothesis), but rather selectively influence control of behavioral programs critical for thermal navigation (setpoint hypothesis).

### Inflammation adjusts the valence of temperature stimuli to induce fever

Inflammation alters specific kinematics of individual swims, suggesting changes to the behavioral algorithm that transforms temperature stimuli into navigation behavior (Figure 4). To identify and quantify the changes in the behavioral algorithm, we used a Markov model. We previously used a similar approach to model thermal navigation and preference seeking ^27^. In this model, behavior is characterized by transitions between three long term behavioral modes ^27^: “Reversals” in which larval zebrafish reverse their direction by stringing together multiple turns (e.g., a fish who is facing one side of the thermal gradient reverses its heading to face the other side), “Persistent swimming” in which they hold direction across multiple consecutive swims, and a “General mode” in which swims are unorganized, leading to frequent changes in direction without reversing. These long term modes are critical to capture temperature preference seeking in larval zebrafish, and a Markov model outperforms models without long term behavioral modes ^27^. Within this “Navigation model” the temperature stimulus controls transition probabilities between the swim modes (“transition model”) as well as the swims generated in each mode (“emission models”). In other words, the transition model determines how the temperature a larva experiences alters transitions between the reversal, persistent and general modes, while the emission model in each mode determines how the kinematics of individual swims (IBI, displacement, turn angle) relate to the stimulus (Figure 5A). For model comparison, we fit a separate Navigation model using Markov-Chain Monte Carlo ^58^ (see Materials and Methods) on data from the PBS injected controls and poly(I:C) injected larvae introduced in Figure 1.

**Figure 5:**
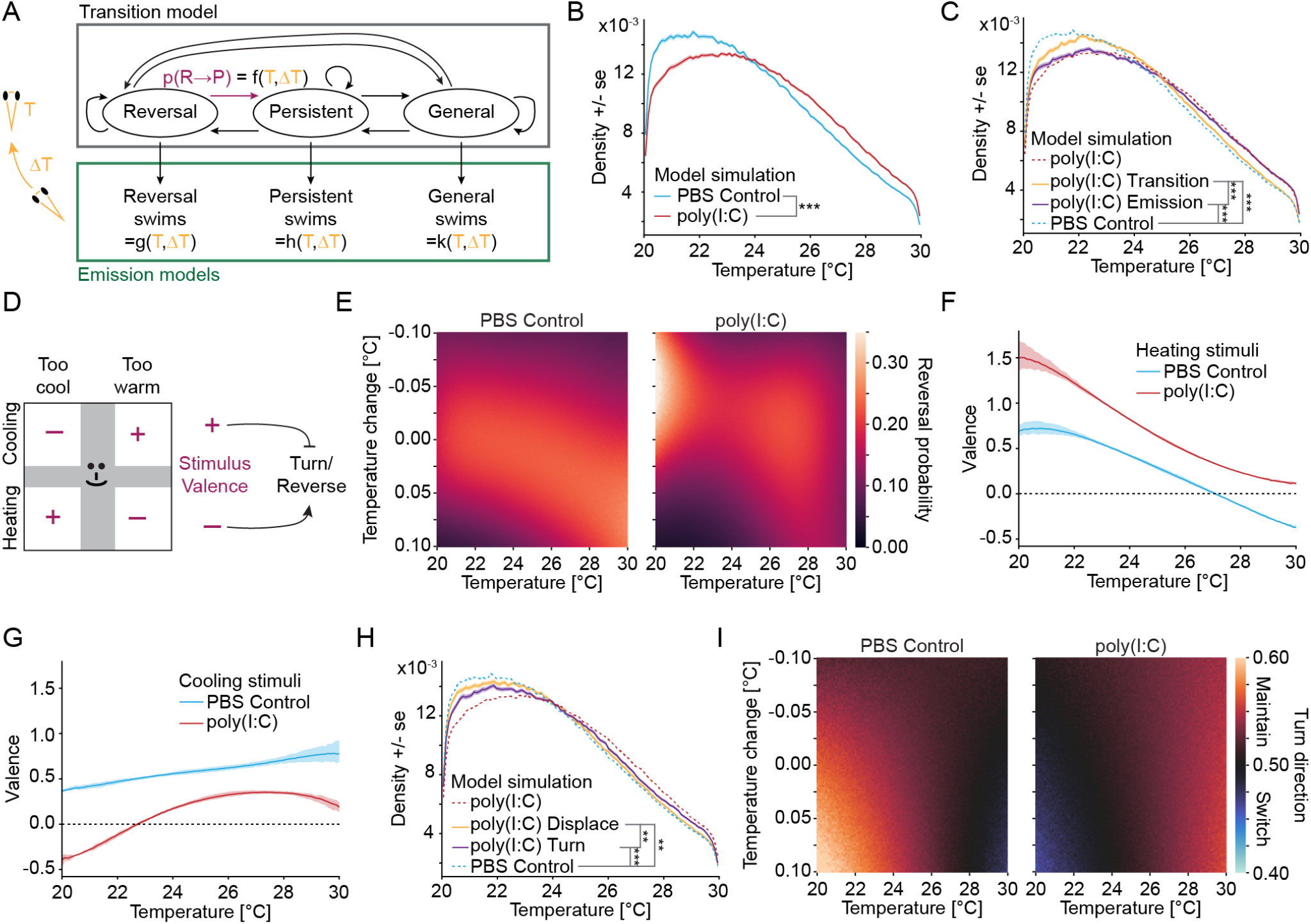
Modeling reveals a switch in stimulus valence underlying fever. **A**: Schematic of the navigation model, consisting of a transition model that encapsulates movements between different behavioral modes (reversals for direction reversals, persistent for swims that keep direction and general for swims that are unorganized), and the mode-dependent emission models that generate the actual swim bouts. Both transitions and emissions depend on the current temperature *T* and the temperature change across the previous swim bout Δ*T*. **B**: Densities of time spent at different temperatures in gradient navigation simulations of models fit on PBS Control data (blue) and poly(I:C) data (red). Shading indicates bootstrap standard error across simulations. **C**: Simulations for hybrid models to test the importance of the transition and emission parts of the navigation model separately. The blue and red dashed lines are replots of the averages shown in F. Simulation of a model with the transition part fit on poly(I:C) data and the emission part fit on control data shown in yellow. Model with the transition part fit on control data and the emission part fit on poly(I:C) data shown in purple. Shading indicates bootstrap standard error across simulations. **D**: Schematic to illustrate important stimulus domains and the relationship between stimulus valence and behavior. Top indicates improving (+) and worsening (-) conditions (positive and negative stimulus valence, respectively) relative to the preferred temperature (vertical gray bar) depending on whether temperatures decrease (cooling) or increase (heating) or are stable (horizontal gray bar). Bottom: since persistent turns and reversal maneuvers indicate changes in swim direction, they are readouts for the valence of the current direction ^27^ which in a gradient is characterized by perceiving heating or cooling stimuli. **E**: Probability of being in the reversal state for the two treatment conditions dependent on the current temperature (X-axis) and the temperature change across the previous bout (Y-axis). The probability is indicated with the color scale depicted to the right. For reference: Bright colors that are restricted to either cooling or heating stimuli indicate reversals that depend on the swim direction within the gradient. **F**: Valence of heating stimuli when experienced at different temperatures for PBS control (blue) and poly(I:C) fish (red). Shading indiates 90% confidence intervals of the posterior distribution. **G**: Same as F but for cooling stimuli. **H**: As in C but comparison of a model in which only the Displacement parts of the emission model have been fit on poly(I:C) data (yellow) with a model in which only the Turn parts of the emission model have been fit on poly(I:C) data (purple). **I**: Turn sequencing in the general swim mode for the two treatment conditions dependent on the current temperature (X-axis) and the temperature change across the previous bout (Y-axis). The color code indicates the probability of the next turn being the same direction as the previous turn - orange shadings indicate that the turn direction is likely maintained (leading to reorientation), blue colors that the turn direction is more likely to switch (no reorientation). In all panels, significance after Benjamini-Hochberg correction is indicated as follows. ***: p-value smaller than critical value at 0.1% FDR; **: p-value smaller than critical value at 1% FDR. See Table S1 for counts and exact p-values. Also see Figure S5

To ensure that comparative modeling would yield useful results, we first tested whether the changes between the Navigation models fit on the different datasets are sufficient to explain behavioral fever. This would indicate that the model replicates the modulation of preference seeking brought about by immune signals. Simulating gradient navigation behavior using the Navigation model (see Materials and Methods) demonstrated that the model encapsulates the changes in the behavioral algorithm underlying fever. Specifically, simulations from the poly(I:C) model sought out significantly warmer temperatures than simulations from the PBS model (Figure 5B). The same can be seen for models fit on the Celebrex data, where poly(I:C) again led to a migration to warmer temperatures, while this effect was abrogated by Celebrex treatment (Figure S5A).

We next used the modular nature of our Navigation model to test whether the transition or emission parts of the model (Figure 5A) were sufficient to drive behavioral fever. To this end, we generated hybrid models in which either the transition or the emission part were derived from poly(I:C) injected data while the other part was derived from PBS control data. This comparison revealed that both changes in the transition and the emission models were necessary for behavioral fever, suggesting changes in both long term behavioral strategies and individual swims. Specifically, both hybrid models sought out significantly warmer temperatures in simulations than a model purely using PBS Control data (Figure 5C). Furthermore, the emission part of the model had a significantly stronger influence on the thermal preference than the transition part (Figure 5C). However, the full poly(I:C) model still sought out significantly warmer temperatures than the model in which only the emission part had been modified (See Table S1) arguing for necessity of the changes to the transition model during fever.

To understand how changes in the transition model give rise to behavioral fever, we analyzed the steady-state probabilities of the individual swim modes and how they relate to the stimulus. A key part of cold- and hot temperature avoidance is the reversal mode. Reversals allow larval zebrafish to efficiently reorient when swimming away from the preferred temperature. They are preferentially triggered during worsening contexts, i.e. times when the fish experiences heating stimuli while above the temperature preference or cooling stimuli while below (Figure 5D). Therefore, we speculated that fever would affect the distribution of reversals within the gradient. To test this, we determined the reversal mode probabilities based on position and swim direction within the gradient from the transition models fit on the PBS injected control data (Figure 5E left) and the poly(I:C) injected behavioral fever data (Figure 5E right). This revealed a clear shift in reversal mode probabilities. In control larvae, reversal probabilities were largely symmetrical at low temperatures, in other words, both heating and cooling stimuli triggered reversals at a similar rate below ∼25 °C. For temperatures above this threshold however, reversal probabilities were higher for increasing than decreasing temperatures, indicating that control fish actively reverse course when swimming towards warmer temperatures (Figure 5E left). This relationship was reversed upon behavioral fever in poly(I:C) injected larvae (Figure 5E right). Here, reversals were more probable for cooling stimuli at temperatures below ∼23 °C while reversal probabilities were largely symmetric at warmer temperatures. In other words, poly(I:C) injected fish actively reversed course when swimming towards cooler temperatures while avoidance of warmer temperatures was reduced. A similar comparison for the probability of being in the persistent or general swim modes (Figures S5B-C) did not reveal major changes, suggesting that differences in long-term behavioral organization were restricted to reversals. Importantly, blocking PGE2 signaling by the addition of Celebrex abrogated the shift in reversal probabilities (Figure S5D) further strengthening the link between behavioral fever and changes in the control of swim modes.

In zebrafish, reversals are avoidance behaviors that are triggered by stimuli with negative valence ^27,28,59^. We therefore inferred the perceived valence of temperature changes in baseline and fever conditions based on reversal probabilities. Specifically, we compared reversal probabilities at different absolute temperatures in cases where temperatures either increase (heating) or decrease (cooling) with the reversal probability when temperatures are stable. An increase in reversal probability for heating over stable temperatures would indicate that the perceived valence of heating stimuli was negative. Behavioral fever triggered a significant change in the valence of both heating and cooling stimuli. Specifically, while heating stimuli had negative valence for temperatures above ∼27 °C in baseline conditions, they had positive valence in fever conditions throughout the tested range (Figure 5F). At all temperatures tested, the valence of heating stimuli was higher in fever than baseline conditions. The opposite was true for cooling stimuli (Figure 5G). At temperatures below ∼23 °C the stimulus valence of cooling stimuli was negative for poly(I:C) injected fish, while it was was significantly higher across the thermal range for PBS injected fish. This strongly suggests that immune signals trigger a change in temperature stimulus valence to induce the fever phenotype. Similar to the changes in individual swim kinematics, the observed changes in valence argue against a reduction in the perception of warmth. If such a reduction would have been the cause of the valence changes, it would be expected that the poly(I:C) valence curves are shifted right compared to the PBS control curves. However, they are instead shifted vertically, arguing that valence did not change because fish experienced temperatures as colder.

Celebrex reversed the changes in valence in agreement with its suppression of behavioral fever (Figures S5E-F). Even compared to untreated PBS controls, Celebrex significantly decreased the valence of heating stimuli at temperatures above ∼27 °C (Figure S5E) while it increased the valence of cooling stimuli in the same range (Figure S5F). Just as the slight reduction in average experienced temperature compared to controls (Figure 3D), this suggests that Celebrex removed baseline inflammation in the larvae which were not kept in a pathogen free state.

Changes in the emission model led to a partial behavioral fever phenotype (Figure 5C), which suggested changes in the generation of individual swim bouts during inflammation. To better understand the nature of these changes, we performed simulations in which individual bout kinematics were controlled by models fit on poly(I:C) data, while the remainder of the model was derived from PBS control data (Figures 5H and S5G). As expected from the lack of influence of fever on interbout intervals (Figure 4B), changing the IBI emission models was not sufficient to induce seeking of warmer temperatures (Figure S5G). Comparing the effect of models in which the generation of either swim displacements or turns were governed by the poly(I:C) data revealed larger effect of turns (Figure 5H). In simulations, these models sought significantly higher temperatures than both PBS control models and displacement models. Similar to full reversals, maintenance of turn direction across successive swims serves reorientation and is used by larval zebrafish to avoid aversive stimuli ^28^. We therefore used the Navigation model to investigate whether turn direction maintenance was affected, specifically in the general mode in which swims are less structured (Figure 5I). We found that in the general swim mode, control fish maintained turn direction during heating stimuli at temperatures below 22 °C. This indicates that when fish left the cold edge of the gradient, they tended to turn back towards it. During behavioral fever on the other hand, poly(I:C) injected fish switched turn direction on these trajectories, effectively going straight and away from the cold edge (Figure 5I). This shows that changes in the perceived stimulus valence also influenced individual behaviors within the general swim mode and not only the control of swim modes themselves.

In summary, we found that immune signals adjust the valence of temperature stimuli, congruent with a hypothesis that inflammation increases the thermoregulatory setpoint. Changes to long term behavioral modes and individual turn sequencing suggest that heating stimuli become more attractive, while cooling stimuli turn aversive. This change in valence likely drives larval zebrafish to seek warmer temperatures and thereby induces behavioral fever.

### Detection of viral patterns adjusts medullary temperature coding

Within a temperature gradient, the valence of temperature changes depends on the location relative to the preference. For example, above the preferred temperature heating stimuli signal worsening conditions, while they signal improving conditions below (Figure 5D). We could previously show that larval zebrafish solve the assignment of valence to temperature stimuli by splitting the encoding of temperature change in the medulla. Specifically, separate pairs of response types encode heating and cooling at warm and cold temperatures, respectively ^27^. Since fever adjusts the valence of temperature changes (Figures 5F-G) we wondered if fever likewise adjusts temperature encoding within the medulla, especially considering that we observed prominent expression of EP3 receptors in proximity to trigeminal sensory axons within the medulla (Figure 2D).

To this end we imaged the medulla of larval zebrafish sixteen hours after PBS (control) injection or poly(I:C) injection (Figure 6A). We segmented the imaging data with Suite2P ^60^ and identified neurons that linearly or nonlinearly encode the temperature stimulus using MINE^61^. Poly(I:C) did not lead to gross reorganizations of neural activity as evidenced by a lack of difference in model correlations (Figures S6A) and a lack of change in indicators of neural response nonlinearities (Figures S6 B-C). As we previously observed, the majority of neurons responded nonlinearly due to rectification caused by the split into neurons that respond during warm versus cold temperatures ^27^. We subsequently used an iterative regression method (see Materials and Methods) to find neurons corresponding to the seven medullary response types we previously identified in both the PBS and poly(I:C) injected fish (Figure 6B). Individual responses revealed high stereotypy within clusters and treatment, but for some response types (e.g., Hot cooling) the encoding seemed to differ between treatments (Figure 6B). There were no significant differences in the numbers of neurons identified for most response types according to treatment, except for Hot cooling neurons, of which there were significantly fewer after poly(I:C) injection (Figure 6C). Notably, all response types were identified in more than half of the imaged larvae, irrespective of treatment (Figure S6D). As we have shown previously, there were more cold-responsive neurons in the zebrafish medulla, than warm responsive ones (CA, C, CC, CH, Figure 6C). Injection of poly(I:C) only had small (albeit significant due to the number of neurons) effects on the responses of neurons that are activated in cold temperatures (Figure S6E-H). However, there were clear and significant differences in warm encoding neurons (Figures 6D-F). Neurons that encode hot temperatures responded with a small delay after poly(I:C) injection (Figure 6D). An even larger effect could be observed for neurons that encode cooling stimuli in warm temperatures. These neurons started to respond at a significantly lower temperature after poly(I:C) injection (Figures 6E and G, HC). Since these neurons signal stimuli with a positive valence, this change may underlie the reduced valence of cooling stimuli after poly(I:C) injection (Figure 5 G). Neurons that encode heating stimuli in warm temperatures only responded at higher temperatures after poly(I:C) injection (Figure 6F). Since these neurons signal negative stimulus valence, this change mirrors the increase in valence for heating stimuli after poly(I:C) injection (Figure 5E). However, while there was an overall significant change in the response of Hot heating neurons (Figure 6F), the change in response threshold was not significant after correcting for multiple comparisons (Figure 6G, HH).

**Figure 6:**
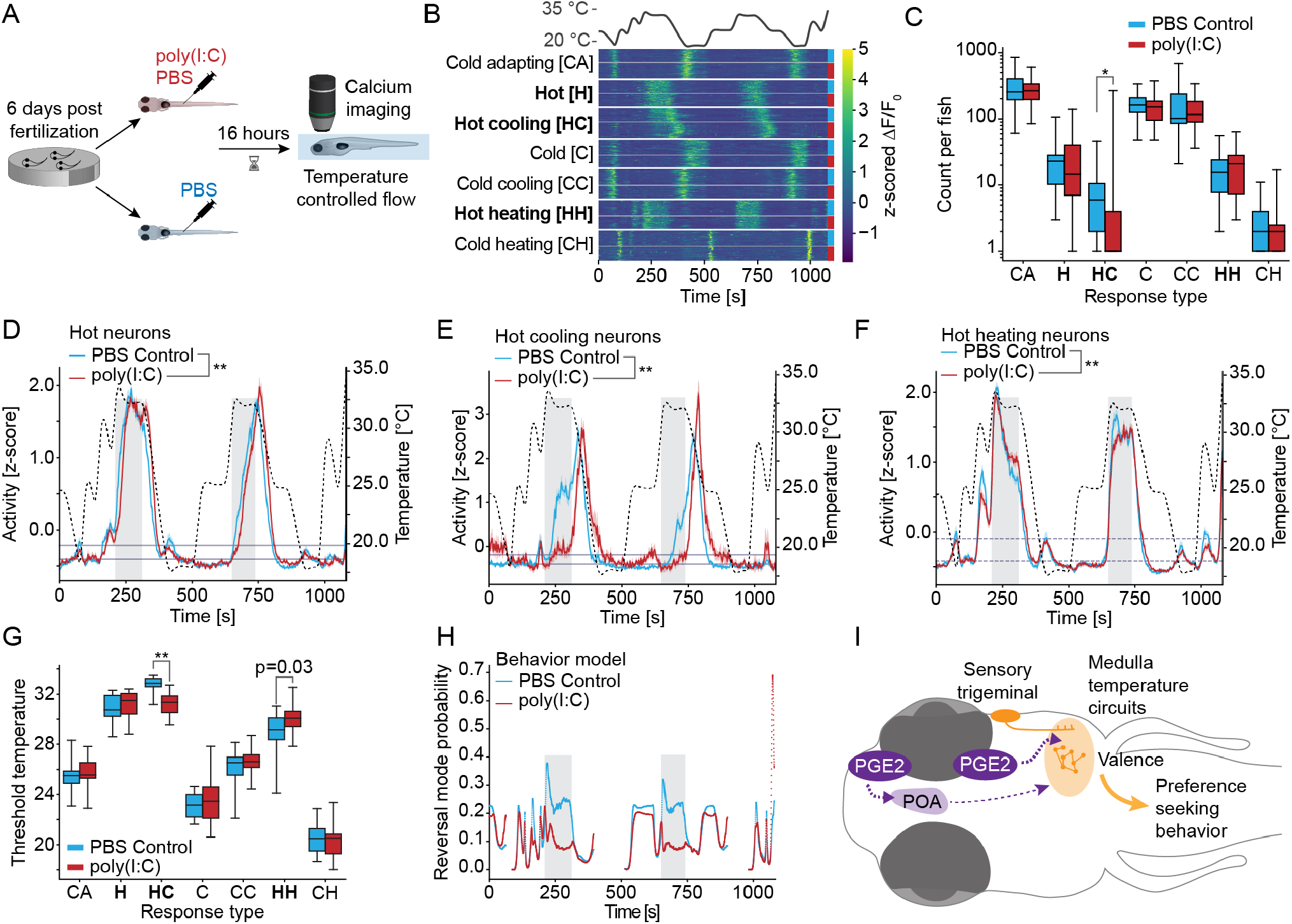
Calcium imaging reveals changes in medullary temperature encoding. **A**: Schematic of the assay and experimental timeline. **B**: Clustered heatmap of randomly selected raw responses across conditions (indicated by colored dots) with temperature stimulus shown on top. Types in bold are those for which traces are shown in D-F. Individual clusters are separated by white lines while thin gray lines separate data from the two treatments within each cluster. Treatments are indicated by the colored bars on the right. The heatmaps color indicates the z-scored fluorescence response of each neuron with the color scale depicted on the right. **C**: For each response type (abbreviations as shown in B) the number of neurons that were identified in each fish that had at least one neuron of the given type according to treatment (PBS Control blue, poly(I:C) red). Mann-Whitney-U test across larvae. **D-F**: Comparison of cluster average responses for response types that respond in warm temperatures. Responses of PBS Control blue, Poly(I:C) red. Temperature stimulus is shown in a black dashed line (right Y-axis). Shaded error indicates bootstrap standard error of responses across larvae. Significance was determined by bootstrapping across the cluster identification procedure (see Materials and Methods). Gray boxes mark the same times in D-F and H. Horizontal navy lines indicate computed baseline and activation thresholds used in G. **D**: Hot response type. **E**: Hot and cooling response type. **F**: Hot and heating response type. **G**: For each type and condition (PBS Control blue, poly(I:C) red) the threshold temperature at which the type activates, based on the thresholds indicated in D-F and Figures S6 E-H. **H**: Reversal mode probability at different times during the imaging experiment. Probabilities are calculated based on the PBS Control navigation model (blue) and the poly(I:C) navigation model (red). Only times in which the stimulus was larger than 20 °C were included since the behavioral model was only fit on those temperatures. Note the times of largest difference between conditions for reversal probability marked by the gray boxes which are replicated in D-F. In all panels, significance after Benjamini-Hochberg correction is indicated as follows. ***: p-value smaller than critical value at 0.1% FDR; **: p-value smaller than critical value at 1% FDR; *: p-value smaller than critical value at 5% FDR. Written out p-values are uncorrected and did not pass criterion after correction. See Table S1 for counts and exact p-values. Also see Figure S6

To better understand the relationship between changes in neural activity and behavioral fever, we used the Navigation models fit on PBS and poly(I:C) data to predict the reversal probability of larval zebrafish during the imaging session (Figure 6H). Since larval zebrafish reduce their swim frequency while embedded within the microscope these data would be difficult to obtain directly. The model prediction suggested two stimulus periods with large changes in reversal probability (gray boxes in Figure 6H). Interestingly, these periods of large difference in reversal probability, also aligned with large differences in neural activity across the three warm response types (gray boxes in Figures 6D-F).

In summary, we show that immune signals alter medullary temperature encoding, especially in response types that signal the valence of heating and cooling stimuli during warm temperatures. This suggests that PGE2 signaling drives fever by modifying temperature valence in the medulla (Figure 6I).

## Discussion

Behavioral fever is the only means of raising body temperature in ectotherms during inflammation ^8,62,63^. However, it is conserved in endotherm animals where it supplements autonomous means of fever induction ^9,10,12–14,21,25,64^. Despite this conservation, we know little about how inflammation drives animals to seek out warmer temperatures, a hallmark of behavioral fever. A hypothesis proposed for both autonomous and behavioral fever is that inflammatory signals directly or indirectly inhibit neurons that signal warmth ^9,14,25,48,65^. This suggests that animals might experience temperatures as colder in febrile conditions. This would then lead to seeking of increased temperatures through a simple subtraction from the temperature measured by sensory neurons in the periphery and the central brain. Alternatively, fever might change the setpoint or preference without altering temperature perception ^66^. However, these hypotheses has not been directly tested, as detailed behavioral and neural measurements of behavioral fever were lacking.

To disambiguate these hypotheses, we analyzed behavioral fever in the vertebrate larval zebrafish. Here, we could show that expression of key genes in cell types implicated in mammalian fever is broadly conserved in zebrafish (Figure 2). Furthermore, viral inflammation leads larval zebrafish to seek out warmer temperatures in a PGE2 dependent manner (Figures 1 and 3), suggesting functional conservation of behavioral fever pathways. Comparing behavioral kinematics in the baseline and fever conditions revealed that larval zebrafish seemingly do not experience temperatures as colder during inflammation, arguing against a global change in temperature perception (Figure 4). Specifically, while temperature dependence of swim displacements and turn angles suggested a shift consistent with a reduction in the perception of warmth, the control of interbout intervals by temperature was unaffected by fever, suggesting that information about “true temperature” is still preserved in fever conditions.

To understand the changes in the behavioral algorithm that lead to seeking of warmer temperatures during inflammation, we performed comparative modeling. Models fit on larvae exhibiting behavioral fever led to a fever phenotype in simulations compared to models fit on larvae in baseline conditions (Figure 5). Comparing the algorithms implemented by the models revealed a shift in the valence of temperature changes, leading to a redistribution of consistent turning and reversal maneuvers. Calcium imaging of temperature encoding neurons in the medulla revealed a specific response shift in neurons that encode temperature change during warm conditions (Figure 6). This change in temperature encoding is congruent with the change in stimulus valence and we therefore propose that PGE2 signaling changes the valence of temperature stimuli to specifically alter behavioral motifs that mediate hot and cold avoidance (Figure 6I). These changes cause larval zebrafish to seek out warmer temperatures, tailoring behavior to the demand of fever without a global change in temperature perception. This strongly suggests that inflammation alters the temperature setpoint rather than changing temperature perception. The conservation of circuits and key molecules involved in fever generation between larval zebrafish and mammals suggests that this change in setpoint and stimulus valence is a conserved principle of fever.

### Localized versus distributed action of immune signals

To induce behavioral fever, inflammatory stimuli must alter processing in thermoregulatory circuits. Rather than a reduction in the perception of warmth, our behavioral and modeling results suggest that PGE2 signaling drives specific behavioral changes by altering the valence of cooling and heating stimuli. These changes could be orchestrated by the detection of inflammatory signals within the preoptic area of the hypothalamus. And we indeed identified neurons that respond to immune signals in mammals ^14,47^ within the POA of larval zebrafish (Figure 2). Preoptic neurons would likely signal through downstream structures such as the hypothalamus and raphe nuclei which could exert neuro-modulatory effects in sensori-motor circuits ^25,67,68^. This circuit organization implies that the POA harbors command-like neurons triggering behavioral and autonomic fever. Importantly, there is clear evidence for an involvement of preoptic neurons in behavioral fever suggesting that the POA serves this role ^14,69^. However, we also identified a prominent cluster of EP3 receptor expressing neurons in the medulla in close proximity to axons of trigeminal sensory neurons (Figure 2). This suggests that PGE2 itself might act as a distributed modulatory signal, altering processing across thermal navigation circuits to induce fever, as opposed to exclusively triggering fever through command-like neurons within the POA. Support for a distributed rather than localized action of PGE2 has also recently emerged in mammals, were EP3 receptor expressing neurons in the medullary parabrachial nucleus mediate warm seeking behavior ^49^. In this framework, immune signals would directly modulate target circuits within the brain to induce an internal fever state. We currently do not know whether direct PGE2 action in the medulla is ultimately responsible for the changes in medullary temperature encoding (Figure 6). However, the delay in responses we observe in medullary neurons during temperature stimulation is consistent with an inhibitory signal from EP3 receptors. Whether immune signals act in a localized or distributed manner to alter the preference also relates to a more general question about the encoding of the preferred temperature setpoint. Specifically, the thermoregulatory setpoint could be encoded in the firing rate of dedicated neurons and alteration to their activity would shift the preference. Alternatively, the preference could emerge as an implicit feature of temperature processing circuits. While our data show that inflammatory signals shift the setpoint rather than the perception of temperature, they cannot disambiguate whether the setpoint is encoded explicitly or is an emergent feature of thermoregulatory processing. However, future studies on distributed or central action of PGE2 signals could shed light on this distinction.

### Neural substrates of temperature valence

The modulation of activity in neurons signaling temperature change within the medulla is consistent with the change in valence inferred from behavioral data. We focused our imaging on the medulla because of the prominent expression of EP3 receptors and our prior work implicating medullary circuits in temperature avoidance and navigation ^26,27^. However, neurons within the right dorsal habenula have been implicated in controlling reversals during heat avoidance in larval zebrafish ^28^ and synaptic changes between neurons of the right dorsal habenula and interpeduncular neurons allow larval zebrafish to associate behavioral actions with successful cold avoidance ^59^. Since the habenula plays a role in mammalian thermoregulation ^52,70^ and is important for signaling stimulus valence ^71,72^, it is likely that part of the changes in stimulus valence we observe are mediated by changes in habenular activity. The dorsal habenula in zebrafish receives inputs from the preoptic area ^73^ and immune signals detected withing the POA could therefore influence habenular activity. Similarly, in mammals the habenula receives preoptic inputs ^74,75^. However, we also observed sparse EP3 receptor expression within the right dorsal habenula in larval zebrafish (Figure S2G). Immune signals could therefore directly reduce habenular activity to alter the valence of temperature stimuli.

## Materials and Methods

Animal handling and experimental procedures were approved by the Ohio State University Institutional Animal Care and Use Committee (IACUC Protocol #: 2019A00000137-R2).

### Fish handling

Larval zebrafish were raised in autoclaved, buffered E3 medium ^76^ with a pH of 7.2 in an incubator with a 14:10 light:dark cycle at 28 °C. All experiments were performed in E3 medium with the addition of drugs and Dimethyl sulfoxide (DMSO) as indicated below. For ZFPV experiments, larval zebrafish were raised specific-pathogen-free. Embryos were bleached as previously described ^35^ with the following modifications: bleach concentration of 46 ppm, 4-minute washes, and a final pronase concentration of 14 *µ*g/mL.

### Larval injection of poly(I:C) or PBS

Microneedles were made using a model p-2000 sutter instrument needle puller set at HEAT=380, FIL=4, VEL=40, DEL=200, PUL=175. 4in thinwall GL 1.2OD/0.9ID glass capillaries from world precision instruments were pulled at this setting. After pulling the capillaries, the tip of the needle was formed using the “bump method”, breaking off the tip until the opening is 6-9 microns across. Injection solution was created by combining 1 *µ*l of 100 mg/ml Rhodamine-dextran dye (Sigma) with either 10 *µ* l of 1 mg/ml high molecular weight poly(I:C) (Invivogen, USA) dissolved in 1x PBS, or 10 *µ*l of 1x PBS for controls.

6 days post fertilization (DPF) zebrafish were anesthetized with 160 *µ*g/ml Tricaine (Syndel, USA) ^76^ and mounted in 2.5 % low melting point agarose (Sigma-Aldrich, USA) at a slight angle, ventral side up to allow for easy access to the caudal vein. The tail was subsequently freed so the needle could access the caudal vein. The glass needles were filled with 5 *µ*l of the injection solution. 10 nl of injection solution was injected into the caudal vein of larval zebrafish. The fish were released from the agarose then left to rest for 16 h at 28 °C until the respective behavioral or calcium imaging experiments.

### ZFPV experiments

ZfPV preparation and administration were adapted from a previous study ^35^. ZfPV preparations were generated from 30 randomly selected adult wild-type Tubingen animals. Animals were euthanized by immersion in ice water, after which intestines were removed and homogenized together in 30 mL of autoclaved E3. The homogenate was subjected to 3 freeze-thaw cycles using liquid nitrogen and a 37 °C water bath. It was then centrifuged at 1, 000 × *g* for 10 minutes, and the resulting supernatant was further centrifuged at 10, 000 × *g* for 30 minutes. The supernatant was passed through a 0.22 *µ*m syringe filter, concentrated using a Vivaspin 20 Centrifugal Concentrator, and stored at -80 °C. The resulting filtrate contained 62,366 ZfPV genomes/*µ*L as determined by RT-qPCR. In experiments using ZfPV, filtrate was applied to 6 DPF or 7 DPF clutch mates for 24 hours via bath immersion at a concentration of 623,660 viral genomes/mL. Filtrate was applied 2 days prior to behavioral experiments.

### Celebrex treatment

In experiments using Celebrex (Tocris, USA), the drug was applied via bath immersion at a 10 [*µ*M] concentration together with DMSO at a concentration of 0.1 % v/v in E3 medium. All control fish for these experiments were incubated in DMSO at a concentration of 0.1 % v/v in E3 medium. After poly(I:C) injection, larvae were immediately transferred into either Celebrex+DMSO or DMSO only. For ZFPV experiments, Celebrex and DMSO were applied 24h before the start of behavior experiments.

### RNA extraction of individual larval zebrafish

RNA was extracted from individual zebrafish using a Picopure kit (Thermo Fisher Scientific, USA). Larvae were euthanized on ice according to the IACUC protocol and individually placed into a tube with 50 *µ*l of Beta-Mercaptoethanol containing extraction buffer (XB) provided by the kit. The larvae were subsequently ground up using a disposable, motorized pestle (VWR, USA). The samples were then frozen at -80 °C for up to 6 months. After thawing on ice, samples were subsequently processed according to the Picopure kit protocol and eluted into RNAse/DNAse free 1.5 ml microcentrifuge tubes (Eppendorf, Germany). The extracted RNA was either used immediately for reverse transcription or stored at -80 °C.

### Reverse transcription reaction

Concentration of RNA was measured using a Nanodrop (Thermo Fisher Scientific, USA). 125 ng of extracted RNA was diluted to 12 *µ*l total volume using ultrapure water. This solution was then pipetted into 50 *µ*l PCR tube with 1 *µ*l of 10 *µ*M dNTPs (Thermo Fisher Scientific, USA) and 1 *µ*l of 50 *µ*M Oligo d(T) (Thermo Fisher Scientific, USA). This solution was incubated for 5 minutes at 65 °C then placed on ice for 1 minute. The sample was processed using Superscript IV RT enzyme according to manufacturer instructions (Thermo Fisher Scientific, USA) and followed by RNAseH digest.

### qPCR on cDNA produced from individual fish

QPCR was performed on a Bio-Rad CFX Connect Thermal Cycler with Optics module (2017) thermocycler using SYBR green master mix (Biorad, USA) according to manufacturer protocol. Each of our qPCR reactions had the same formula; 10 *µ*L of undiluted SYBR green master mix; 0.5 *µ*L of forward primer;0.5 *µ*L of reverse primer;4 *µ*L of cDNA reaction diluted 1:3 with H2O; 5 *µ*L of H2O. 58 °C was chosen as the annealing temperature. All tested transcripts were normalized to Mob-4. See Table S2 for primer sequences.

### HCR In situ hybridization

HCR *in situ* hybridization was performed according to a mixture of manufacturer protocols (Molecular Instruments, USA) and previously published variations as described below ^77,78^. Probes were produced by Molecular Instruments and buffers indicated below are the corresponding buffers of the Molecular Instruments kit. 7 DPF larval zebrafish were euthanized on ice according to the IACUC protocol then fixed overnight in 4 % Paraformaldehyde (PFA) in 1x PBS at 4 °C on a nutating mixer. The next day, fish were given a quick rinse then washed three times for 5 minutes in 1x Dulbecco’s Phosphate buffered saline shaking at 150 rpm. Larvae were subsequently permeabilized for 10 minutes (strict timing) in 100 % (v/v) pre-chilled methanol at -20 °C. Larvae were rehydrated in 1:1 Methanol/2x SSCT (v/v) for 5 minutes on ice. Two additional washes of 2x SSCT for 5 minutes at room temperature (RT) were performed while shaking at 200 rpm. The samples were then placed in 500 *µ*l of pre-warmed (37 °C) hybridization buffer for 30 minutes. 500 *µ*l of pre-warmed (37 °C) hybridization buffer was then combined with 1 *µ*M probes; 2uL of Gad1b probe was used and 4ul of probe was used for ptger3, ptger2b, and galanin since they had low expression levels. This solution was then added to the fixed and washed larval zebrafish and left to hybridize overnight at 37 °C while shaking at 200 rpm. The next day, the samples were washed with 500 *µ*l of pre-warmed wash buffer for 20 minutes. The samples were subsequently washed twice with 2X SSCT for 20 minutes at RT. The larval zebrafish were left to pre-amplify in amplification buffer at RT for 30 minutes. Hairpins (H1 & H2) were linearized at 95 °C for 1 minute then placed in a dark place to rest for up to an hour at RT. Amplification buffer was created by combining 100 *µ*l of amplification buffer with 4 *µ*l of 2 *µ*M probe. This solution was then added to the samples drained of the pre-amplification buffer and left at RT overnight, shaking at 150 rpm. The next day, the samples were washed three times in 2x SSCT for 20 minutes at RT under foil. The samples were subsequently imaged under a custom-built 2-photon microscope.

#### HCR image acquisition and registration

After staining, image stacks were acquired on the same custom-built 2-photon microscope used for calcium imaging (see below). For each probe as well as amplifier-only controls, stacks for the anterior half of the brain (forebrain and parts of the midbrain) and the posterior half (parts of the midbrain and the hindbrain) were acquired separately with a pixel-size of 0.65 *µ* m across planes spanning the entire dorso-ventral extent at 2 *µ*m step size. For each larva, these anterior and posterior stacks were registered separately to a previously generated reference brain ^26^ using non-rigid image registration through CMTK^79^. The anterior and posterior stacks were merged after registration. Since there was a large zone of overlap between the anterior and posterior stacks, the stacks were merged such that each voxel in the final stack only had contributions from one of the two stacks. Small artifacts are visible at the merge line, however this approach was chosen over stitching since it was more precise than stitching the stacks before registration while stitching post-registration led to changes relative to the common coordinate system of the reference. Images displayed in Figures 2 and S2 are maximum intensity projections after averaging three larvae.

### Behavioral assays

Behavioral assays were performed in a previously described gradient chamber ^27^. All experiments were performed in a gradient from 20 °C on the cold to 30 °C on the hot side. Since the gradient chamber has two lanes, in each experiment two larvae were assayed, one per lane. However, to avoid possible confounds of likely small differences in temperature profiles between the lanes, in each experiment two larvae of the same treatment were assayed. To further avoid potential confounds related to assay timing, care was taken that the test order of treatments was shuffled between experimental days. On each experimental day, larvae from all treatment conditions were tested. Before each experiment, the chamber was allowed to stabilize and the gradient was checked with a thermal camera (Fluke, USA). Each experiment had a duration of 30 minutes and the first 5 minutes of experimental time were excluded from analysis to account for possible handling artifacts. Positions and swim kinematics were extracted and analyzed as described previously ^27^. In brief, larval positions and heading angles were extracted online during the experiments. These were subsequently used to extract bouts and subsequently interbout intervals (IBI), swim displacements and turn angles as well as turn magnitudes (absolute value of the turn angle). These kinematic quantities were analyzed by temperature while the positions themselves were analyzed to determine occupancy of the larvae within the gradient. As before, swim bouts within 4 mm (one larval length) of any edge of the chamber were excluded from all further analysis to avoid confounds by the edges influencing swim direction and turning. To analyze the effect of bout temperature on swim kinematics, weighted histograms (binned according to temperature) were calculated in each individual larva and subsequently averaged across individuals.

### Navigation model fitting and analysis

Swim trajectories and swim modes (Reversal, Persistent, and General) were assigned as previously described based on swim kinematics across groups of adjacent swim bouts ^27^. In brief, trajectories consisted of swim bouts in continuous time slices unbroken by edge excluded bouts or tracking errors. Within trajectories, swim modes were assigned to individual swim bouts based on whether they were part of direction reversals (“Reversal mode”), continuous directional swims (“Persistent mode”) or neither (“General mode”) ^27^.

All models were fitted using Hamiltonian Markov chain Monte Carlo via STAN^80^. For model-fitting, the respective design matrices *X* were orthogonalized using QR-Decomposition within STAN. Model parameters *θ* were fit in this space and subsequently put back into the original space using the inverse transform. Linear dependencies were thus removed from the fit while still obtaining parameters *β* that could be used on raw data during tests and simulations.

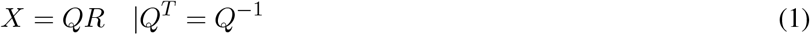

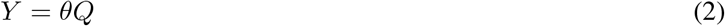

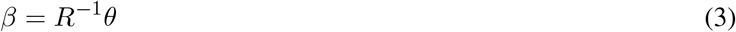

Swim bouts were grouped into trajectories and 80 % of all trajectories were used to fit models (Markov model transitions, emission models) while the remaining 20 % were held out for model testing (data not shown). Trajectories were split into the train/test set by adding 4 consecutive trajectories into the train set, followed by adding one trajectory into the test set and so on. Therefore, each set was made up of trajectories across all fish. All presented model data (e.g., steady-state probabilities, simulations) are based on individual model draws rather than a maximum likelihood average. For aggregate data such as stead-state probabilities, averages across 1000 such draws are presented in the figures. For simulations, a different random draw of model parameters was used at each timestep.

#### Markov model of swim modes

The Markov model was defined to capture transitions between the three swim modes, reversal mode (r), persistent mode (p) and general mode (g). The transition probabilities between modes *M* were set by a Generalized Linear Model (GLM) with the current temperature, temperature change across the previous bout and their interactions as inputs such that *p*(*M*_*t−*1_ → *M*_*t*_) = *f* (*T*, Δ*T*). For the temperature input *T* to the model, 25 °C was subtracted, to roughly center the input. In the stimulus driven model, the log probability of one of the three transitions from each swim mode was pinned to zero (to ensure identifiability of the resulting model), leading to the following definition of transition probabilities:

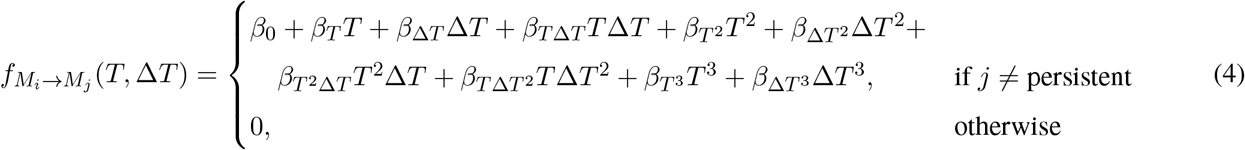

The 3rd order polynomial relationship above was chosen to allow for shifts in the preferred temperature between conditions which would have been harder to parametrize in the original formulation used in *Balakrishnan & Haesemeyer* ^27^Transition probabilities were then computed using the softmax function:

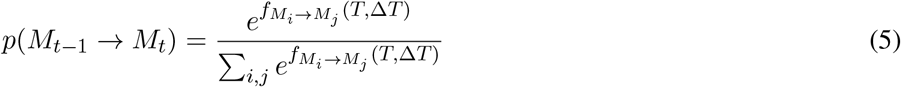

#### Emission models of swim bout kinematics

To capture swim bout generation dependent on swim mode as well as the stimulus and behavioral history the following emission models were fit. For each model, the training data was first split according to swim mode and subsequently separate emission models were fit for each mode. The rationale for this approach was that modes were likely differentiated by eliciting different swim bouts. At the same time the expectation was that within each swim mode, bout kinematics would be influenced by the stimulus as well. Hence separate GLMs relating stimulus features to kinematic distributions were fit for each mode. As for the transition model above, functional representations were chosen that allow to encapsulate arbitrary preferences.

##### Turns

Turns were modeled as a Gaussian-Gamma mixture model. Straight swims were captured using a Gaussian distribution centered at 0 with a standard deviation *σ*_*s*_ fit on the data. Left and right turns were modeled as mirror-symmetric Gamma distributions with parameters shape *α*_*ϕ*_ and rate *β*_*ϕ*_ fit on the data. *σ*_*s*_, *α*_*ϕ*_, *β*_*ϕ*_ were fixed for all turns within each mode but were fit independently for each mode. Within each mode, turn distributions were modeled as mixtures of these distributions with the mixing coefficients Θ dependent on the current temperature *T*, temperature change Δ*T* of the previous bout as well as the previous turn angle *ϕ*_*t−*1_, according to:

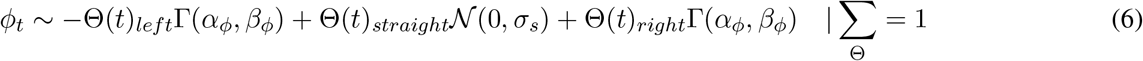

with

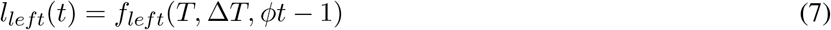

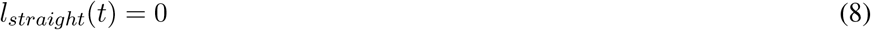

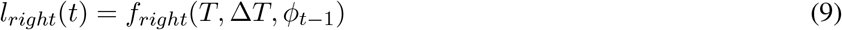

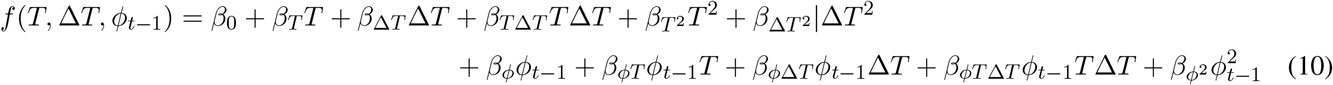

and

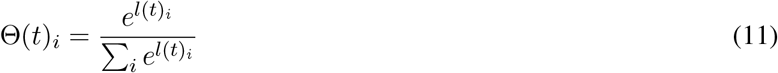

##### Displacements

Displacements *d* were modeled as a single Gamma distribution since displacements are strictly positive. While not a perfect description of displacements, out of the standard distributions displacements were best fit with this distribution. The distribution was parameterized using rate *β* and average *µ* such that:

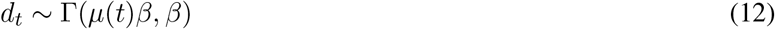

This parameterization was chosen under the assumption that larval zebrafish control the average displacement. It also yielded more stable fits than a standard parameterization according to shape and rate. The rate parameter *β* was fixed for each swim mode while *µ*(*t*) was modeled as a function of the current temperature *T*, temperature change Δ*T* of the previous bout as well as the previous displacement *d*_*t−*1_, according to:

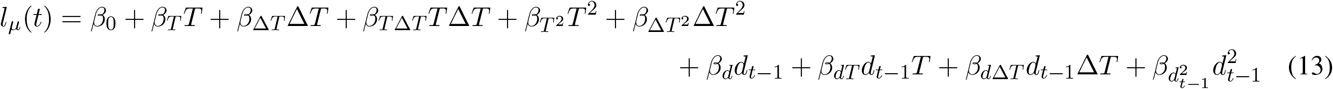

and

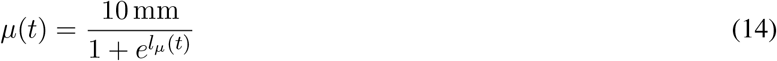

In this formulation 10 mm was chosen as the maximal average displacement. Given that average displacements irrespective of the stimulus never exceeded 4 mm this was a mild constraint on the fit.

##### Interbout intervals

Interbout intervals were well described as Gamma distributions and fit in the same way as bout displacements with the only difference being that the average bout displacement was modeled as a function of temperature and temperature change only, since auto correlations were low for inter-bout intervals. Therefore:

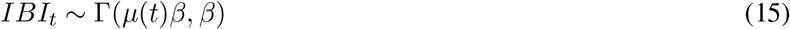

with

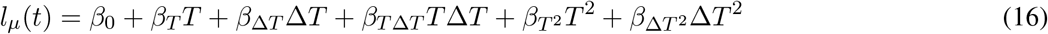

and

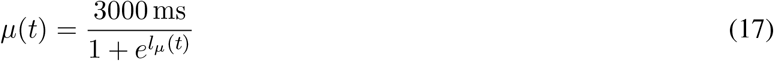

#### Steady state probabilities, valence, and turn maintenance

Steady state probabilities *p*(*M*) for each swim mode *M* for different inputs were computed by exponentiating the transition matrix Θ 100 times. This approach was chosen for efficiency over repeated eigen-decomposition, however we note that the steady state probabilities can be computed analytically as the right eigenvector with the highest eigenvalue (1) of the transition matrix. These approaches are equivalent as long as the transition matrix does not describe an oscillatory system.

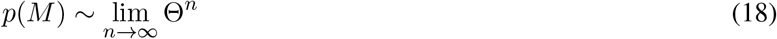

For each tested combination of *T* and Δ*T* steady-state probabilities were computed across 1000 draws from the posterior distribution of parameters and average values are reported in Figures 5 and S5.

The steady state probabilities of reversals *p*(*M*)_*r*_ were used to infer valence as reversals are strong indicators of stimuli that signal worsening conditions to larval zebrafish ^27^. The temperature change Δ*T* across the previous swim bout was used to classify swims into those that are in direction of increasing temperatures (“heating”) for Δ*T >* 0.025, decreasing temperatures (“cooling”) for Δ*T* < −0.025, or “no change” for *−*0.025 ≤ Δ*T* ≤ 0.025. Using the same steady-state probabilities displayed in Figures 5 and S5 the valence of heating (cooling) stimuli was subsequently defined as the difference in average reversal probability between the no change condition and heating (cooling) conditions. A positive valence therefore indicates that reversals are less likely during heating (cooling) than when temperatures do not change.

The probabilities to maintain turn direction (*p*_*maintain*_) in the general mode according to *T* and Δ*T* were computed using - (10). Specifically,

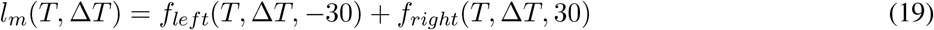

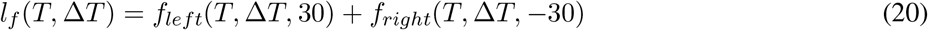

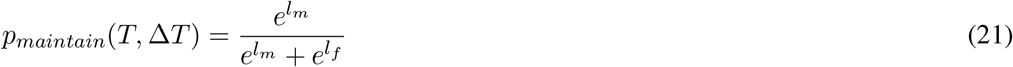

As for the steady-state probabilities, individual draws from the posterior distribution of parameters were used and averages across draws are reported in Figure 5.

### Navigation model simulations

Since all analysis of the fish in experiments assumed the fish as a point particle with a defined centroid and heading angle, the virtual fish in simulations was modeled as a point particle with an associated vector as a heading direction as well. A virtual chamber was defined according to identical dimensions as the experimental setup, with a 200 mm length and 50 mm width. The particle was started at a random point in this chamber with a random heading direction, and was moved along the chamber by generating bouts according to the Navigation model (Figure 5A) with stimulus-controlled. Each simulation was run for a length of 10 hours (3.6 million frames) at 100 Hz to obtain steady-state conditions. As in the experiments, a gradient ranging from 20 °C to 30 °C was simulated. Edge conditions were resolved such that whenever a movement would take the particle across the edge of the virtual arena, its direction of movement was inverted, i.e. the fish was reflected away from the edge. This approach does not mimic edge-tracking behavior (thigmotaxis) observed in larval zebrafish, but since no comparisons between simulated data and behavioral data were performed, this does not pose an issue. Importantly, removing the simulated particle from the edge allowed increasing the amount of data generated in regions of the chamber that matter for comparisons across simulations.

At the start of each simulation the particle was set to be in the General swim mode. Subsequently swim bouts were chosen in the following manner. The Markov model was used to implement a probabilistic state transition with the transition matrix being computed using the current temperature and the temperature change across the previous swim bout. For computation a random parameter sample was drawn from the posterior distribution. After the state transition, a turn angle, displacement, and interbout interval were drawn according to the probability distributions generated by the emission models for the newly selected state. Characteristic parameters of the distributions (see: Emission models) were computed using temperature, temperature change, previous bout angle, and previous displacement by drawing a random parameter sample from the swim modes posterior distributions. Subsequently a random turn angle, displacement, and interbout interval were drawn from the emission distributions. The angle and displacement were used to implement movement of the particle while the interbout interval was used as a waiting time until the next swim mode transition and bout implementation.

While the navigation model consists of a transition model and emission models for each swim mode (see above), all these sub-models are independently fit and used sequentially during simulations. It is therefore possible to generate hybrid models, which were utilized as follows. The mixed model simulations were performed by combining parts of the model fit on PBS injected control data with parts of the model fit on poly(I:C) injected data (experiments from Figure 1. For figure 5G, “poly(I:C) Transition”, transition probabilities were drawn according to the poly(I:C) model, while emissions were drawn according to the emissions models fit on the PBS data. For the “poly(I:C) Emission” model this was reversed and transition probabilities were drawn according to the PBS injected data while emissions models fit on the poly(I:C) injected data were used. For figure 5H, all transition probabilities and emissions were drawn according to the PBS injected data except either the displacement emissions (“poly(I:C) Displace”) or the turn emissions (“poly(I:C) Turn”) which were drawn from the poly(I:C) injected data.

### Calcium imaging during temperature stimulation

Functional calcium imaging in the medulla (hindbrain) was performed on a custom-built two-photon microscope ^61^ at 2.5 Hz with a pixel resolution of 0.78 *µ*m*/*pixel and at an average power of 12 mW at sample at 950 nm (measured optical resolution of the system is *<* 1 *µ*m lateral and *<* 4 *µ*m axial). Larval zebrafish were mounted in a custom designed chamber in which water was flowing through a channel. During imaging the head of the larva was fixed in 2.5 % medium melt agarose (Fisher Scientific, USA) while the tail was kept free to move. To generate different flow temperatures, an inline flow heater (Warner Instruments, USA) with associated controller was used. Control temperatures were calibrated such that at sample temperatures in the range from 18 °C to 32 °C were reached. Temperatures at sample were continuously monitored through a thermistor (Warner Instruments, USA) placed behind the larva and were recorded during the experiment. During imaging, scan stabilization along the z-axis was used to avoid artifacts induced with heating and cooling the agarose with the flowing water ^26^.

For temperature stimulation, within each imaging plane a 18 minutes long stimulus was presented to the larval zebrafish. The stimulus was the same as we used previously ^27^ to allow for the activity clustering described below. Imaging planes were spaced 5 *µ*m apart and 10-15 planes were imaged in each fish.

On each experimental day, one PBS injected control larva and one poly(I:C) injected larva were tested. The order of imaging was randomized across days.

### Processing of 2-photon data

Raw imaging data was processed using Suite2P ^60^ to extract individual neural responses in each plane. Suite-2P was finetuned to properly identify neurons in our imaging setup. Subsequently MINE was used to identify neural responses that could be predicted (either linearly or nonlinearly) based on the temperature stimulus ^61^. A correlation cutoff on test data of *r* = 0.6 was used to decide that a neuron’s activity was related to the stimulus and all neurons exceeding this cutoff were used for the subsequent analyses. The distributions of test data correlations and nonlinearity metrics in Figure S6 were directly derived using MINE.

#### Response type identification across poly(I:C) and control fish

We previously identified seven neural temperature response types within the zebrafish medulla ^27^. To identify these response types within the current datasets while allowing for changes to temperature encoding, an iterative convergence method was used. The cluster average responses obtained on the previous dataset ^27^ were used as initial regressors ^81^ to identify candidate neurons. For each response type regressor all neurons within the poly(I:C) or control dataset with a correlation 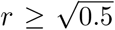 were considered response type candidates. All candidate neurons for each type were averaged to obtain the regressors for the next iteration. These averages were subsequently used to identify the next set of candidates based on the same correlation threshold. Iterations were stopped once successive averages had converged such that the correlation of successive averages was *r >* 0.99. At each iteration, if a candidate neuron was correlated to more than one response type above threshold it was assigned to the response type with the higher correlation.

The overall rationale for the approach was that if treatment changed the response, it would lead to a shift in the average encoding that is small enough to still have outliers within the correlation neighborhood of the original response.

### Analysis of single-cell RNA sequencing data

We downloaded the raw 10x Genomics data belonging to the Shafer dataset from the gene expression omnibus. We used cellranger 7.0.1 to count transcripts and prepare it for export to the Seurat V4, package in R^82^. We used default parameters in all analysis steps; Principal components for RunPCA=10; Dimensions for RunUMAP=1:10; Dimensions for FindNeighbors=1:10; Resolution for FindClusters=1. Data was scaled and normalized then FindAllMarkers was used to find markers of each cluster. SCType was used to subcluster all post mitotic neurons based on the positive marker Elavl3, and the negative marker, MKI67. After subsetting, the cells assigned “neurons” by SCType were transformed using SCTransform, RunPCA, RunUMAP, FindNeighbors, FindClusters using the same default setting as before. After another scaling and normalization, cell types were once again assigned using SCType. This time, the default SCType cell type marker file was adjusted to contain orthologous zebrafish transcripts and used.

### Statistics

Non-parametric statistics were used to analyze all data since it is unlikely that any data arose from normal distributions. Whenever more than two conditions were present, statistics on all combinations were evaluated and significance was determined using the Benjamini-Hochberg method. In some cases, raw p-values were presented within figures to illustrate effects as indicated in the respective figure legends.

Error shadings in linegraphs were reported as bootstrap standard errors across larvae (for experimental data) or simulations (for model simulations). For gradient occupancy, a bootstrapped version of the Kolmogorov-Smirnoff (KS) test was used (see *Anderson, et al*. ^30^) to allow computing the statistic across larvae/simulations where most of the variability arises instead of computing it across individual timepoints. To test for significant differences in the effect of temperature on bout kinematics (e.g., Figure 1F) a bootstrap hypothesis test was used comparing the mean squared distance across temperature bins across fish. This statistic reflects an overall effect rather than testing signficant differences in individual temperature bins. The same statistic was used to compare neural activity across conditions in Figure 6 and S6. However, since the variability here is largely driven by the clustering approach, instead of bootstrapping the statistic across larvae, it was bootstrapped across repeated runs of the clustering algorithm on subsets of the data drawn with replacement.

## Acknowledgements

Research reported in this publication was supported by the National Institute Of Neurological Disorders And Stroke of the National Institutes of Health under Award Number R01NS123887 to MH, by the Office Of The Director of the National Institutes of Health under Award Number R24OD037693 to MH, the National Institute of General Medical Sciences of the National Institutes of Health under Award Number R35GM142950 to JAG, the National Institute of Allergy and Infectious Diseases of the National Institutes of Health under Award Number T32AI055434 to CET, and a seed grant from the Office of the Vice President for Research at the University of Utah to JAG. The funders had no role in study design, data collection and analysis, decision to publish, or preparation of the manuscript. All authors received salary from the funders. The content is solely the responsibility of the authors and does not necessarily represent the official views of the National Institutes of Health.

We thank Inbal Shainer for invaluable advice on the HCR protocol.

We thank Jessica Osterhout, Lindsay Anderson, and Danica Matovic for comments and suggestions on the manuscript.

## Author Contributions

Conceptualization: M.H.

Investigation: B.C., C.E.T., H.L.H

Methodology: B.C., C.E.T., H.L.H, J.A.G., M.H.

Data curation: B.C. and M.H.

Formal analysis: M.H., B.C., C.E.T.

Funding acquisition: M.H., C.E.T, J.A.G.

Software: K.A.B., B.C., M.H.

Supervision: M.H., J.A.G.

Writing - original draft creation: B.C., M.H.

Writing - review and editing: B.C., C.E.T., H.L.H, J.A.G., M.H.

## Supplemental Figures and Tables

**Table S1**: Details of statistics across all experiments.

**Table S2**: Q-PCR primer sequences.

**Table S3**: Official zebrafish gene names for common names used in Figures 2 and S2.

**Figure S1:**
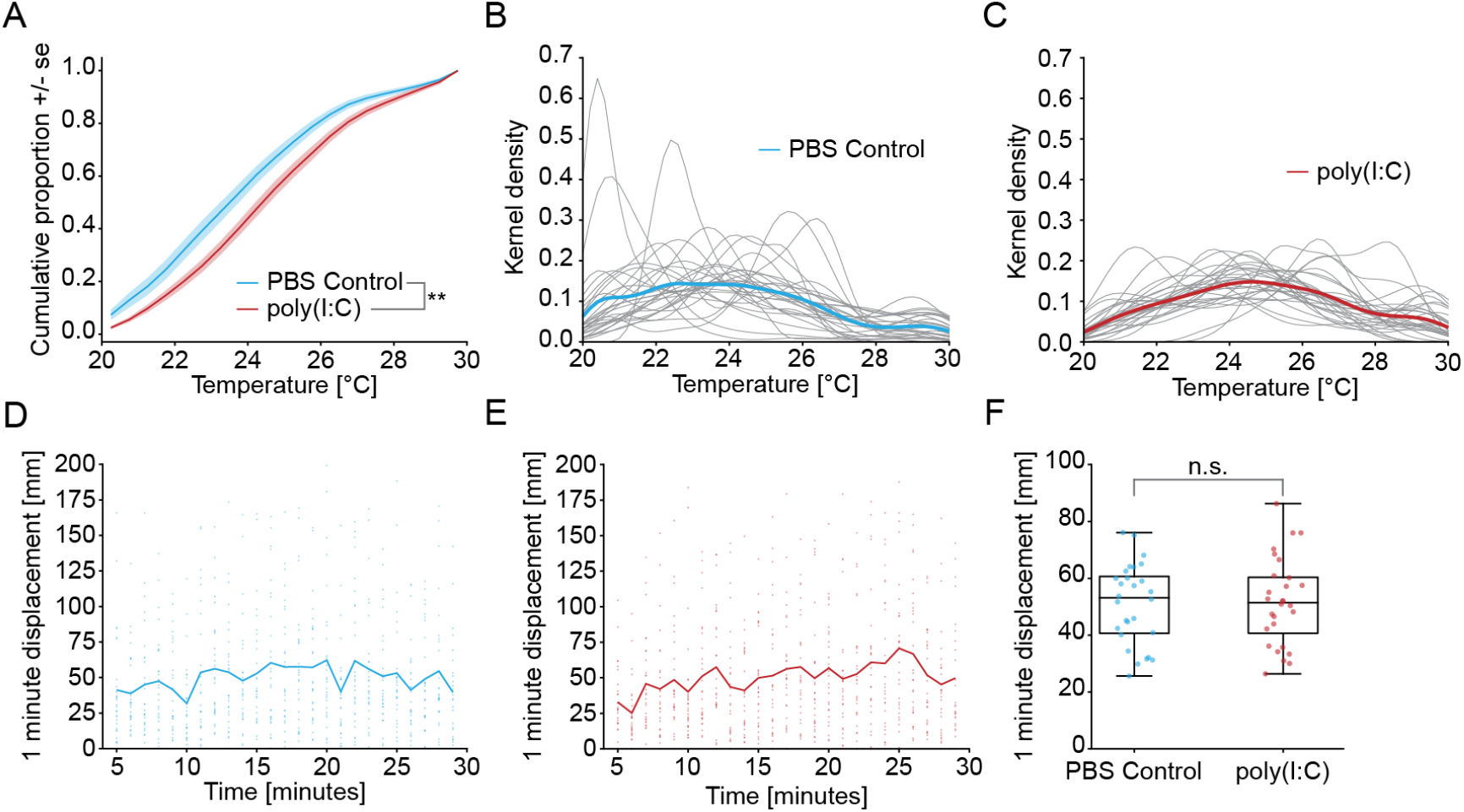
Related to Figure 1. **A**: Cumulative distribution of time spent at different temperatures within the gradient for PBS Control fish (blue) and poly(I:C) fish (red). ∗ ∗ : *p <* 0.01 bootstrapped KS-test across fish. **B-C** Kernel density estimates (KDE) of individual fish distributions within the gradient (grey). Thick colored lines indicate the average across all kernel densities. **B**: Individual PBS injected control fish. **C**: Individual poly(I:C) injected fish. **D-E**: Displacement of the end point of a one-minute long trajectory from the start point of the trajectory for every experimental minute in individual fish (dots) and average (line). Note that the value of this displacement depends both on how vigorously the larva swam and how curved or straight the trajectory was. As such it quantifies the exploratory radius of the fish. **D**: PBS injected control fish. **E**: poly(I:C) injected fish. **F**: Comparison of experiment-average 1-minute displacements in individual fish across treatment conditions. *n*.*s*. : *p >* 0.05 Mann-Whitney-U test across fish.

**Figure S2:**
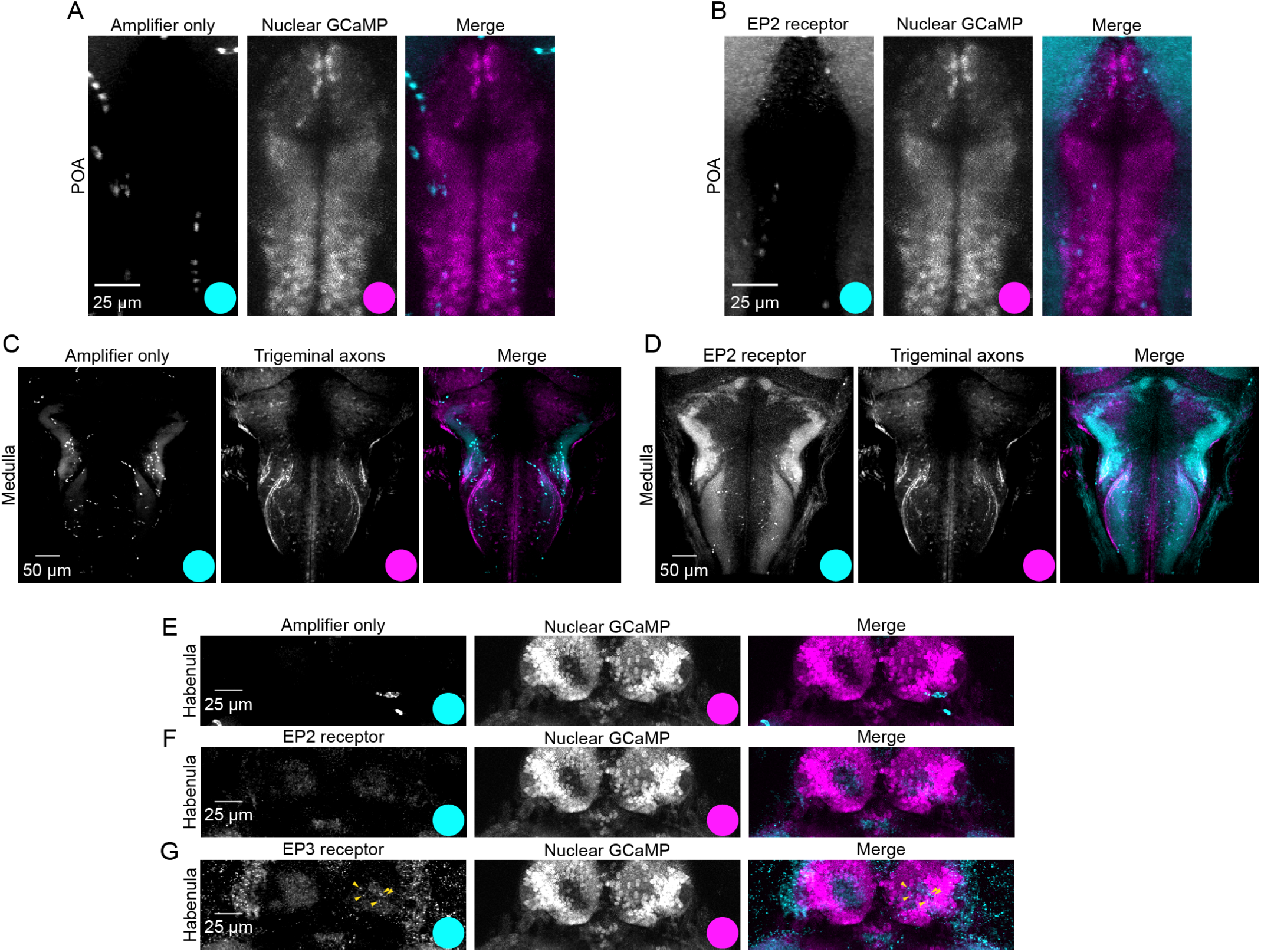
Related to Figure 2. **A**: Amplifier only control of HCR *in situ* in POA neurons. Left panel: Signal of amplifier without probe averaged across 3 registered brains, middle expression of Elavl3-H2B:GCaMP6s marking neuronal nuclei, right panel is merge of both channels. Colored dots indicate the color assigned to each signal in the merge. **B**: Same as A but signal of EP2 receptor probe. **C**: Amplifier only control of HCR *in situ* in the medulla. Left panel: Signal of amplifier without probe averaged across 3 registered brains, middle photo-activatable GFP labeling of trigeminal ganglion sensory neurons revealing location of their projections, right panel is merge of both channels. Colored dots indicate the color assigned to each signal in the merge. **D**: Same as C but signal of EP2 receptor probe. **E**: Amplifier only control of HCR *in situ* in dorsal habenula neurons. Left panel: Signal of amplifier without probe averaged across 3 registered brains, middle expression of Elavl3-H2B:GCaMP6s marking neuronal nuclei, right panel is merge of both channels. Colored dots indicate the color assigned to each signal in the merge. **F**: Same as E but for EP2 receptor probe. **G**: Same as F but for EP3 receptor probe. Yellow arrowheads indicate neuronal EP3 receptor expression.

**Figure S3:**
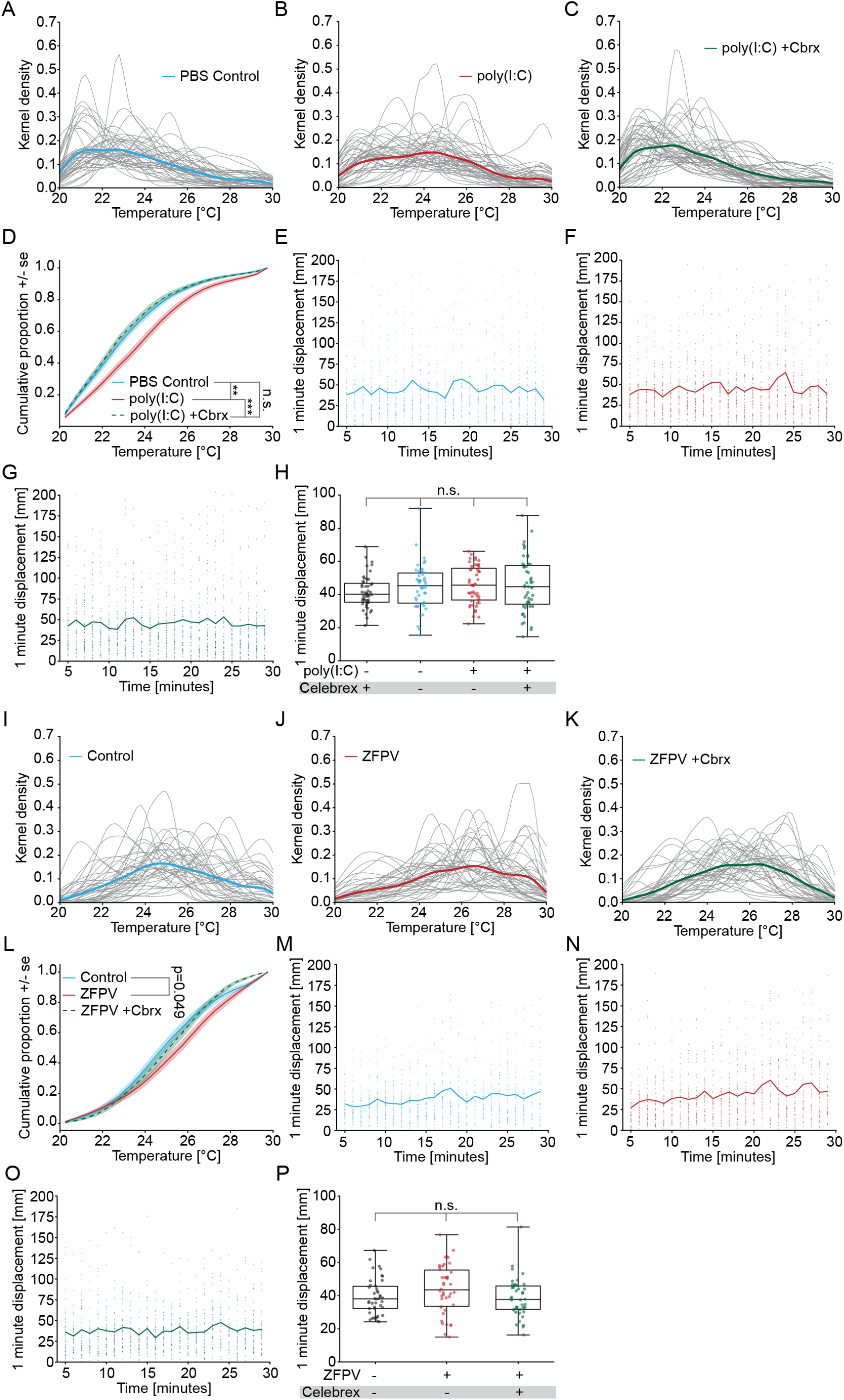
Related to Figure 3. **A-C** Kernel density estimates (KDE) of individual fish distributions within the gradient for Celebrex experiments (grey). Thick colored lines indicate the average across all kernel densities. **A**: PBS injected control fish in DMSO. **B**: poly(I:C) injected fish in DMSO. **C**: poly(I:C) injected fish in DMSO and Celebrex. **D**: Cumulative distribution of time spent at different temperatures within the gradient for PBS Control larvae in DMSO (blue), poly(I:C) fish in DMSO (red), and poly(I:C) fish in Celebrex (dashed green). Shading indicates bootstrap standard error across larvae. Bootstrapped KS-test across fish. **E-G** Displacement of the end point of a one-minute long trajectory from the start point of the trajectory for every experimental minute in individual fish (dots) and average (line). Note that the value of this displacement depends both on how vigorously the fish swam and how curved or straight the trajectory was. As such it quantifies the exploratory radius of the fish. **E**: BS injected control fish in DMSO. **F**: poly(I:C) injected fish in DMSO. **G**: poly(I:C) injected fish in DMSO and Celebrex. **H**: Comparison of experiment-average 1-minute displacements in individual fish across treatment conditions. Mann-Whitney-U test across larvae. **I-K**: Same as A-C but for ZFPV experiments. **I**: Germ free control fish in DMSO. **J**: ZFPV infected fish in DMSO. **K**: ZFPV infected fish in DMSO and Celebrex. **L**: Cumulative distribution of time spent at different temperatures within the gradient for Germ-free control larvae in DMSO (blue), ZFPV infected larvae in DMSO (red), and ZFPV infected larvae in Celebrex (dashed green). Shading indicates bootstrap standard error across larvae. Bootstrapped KS-test across larvae. **M-O**: Same as E-G but for ZFPV experiments. **M**: Germ free control larvae in DMSO. **N**: ZFPV infected larvae in DMSO. **O**: ZFPV infected larvae in DMSO and Celebrex. **P**: Comparison of experiment-average 1-minute displacements in individual fish across treatment conditions. Mann-Whitney-U test across larvae. In all panels, significance after Benjamini-Hochberg correction is indicated as follows. ***: p-value smaller than critical value at 0.1% FDR; **: p-value smaller than critical value at 1% FDR; *: p-value smaller than critical value at 5% FDR; n.s: p-value larger than critical value at 5% FDR. Written out p-values are uncorrected and did not pass criterion after correction.

**Figure S4:**
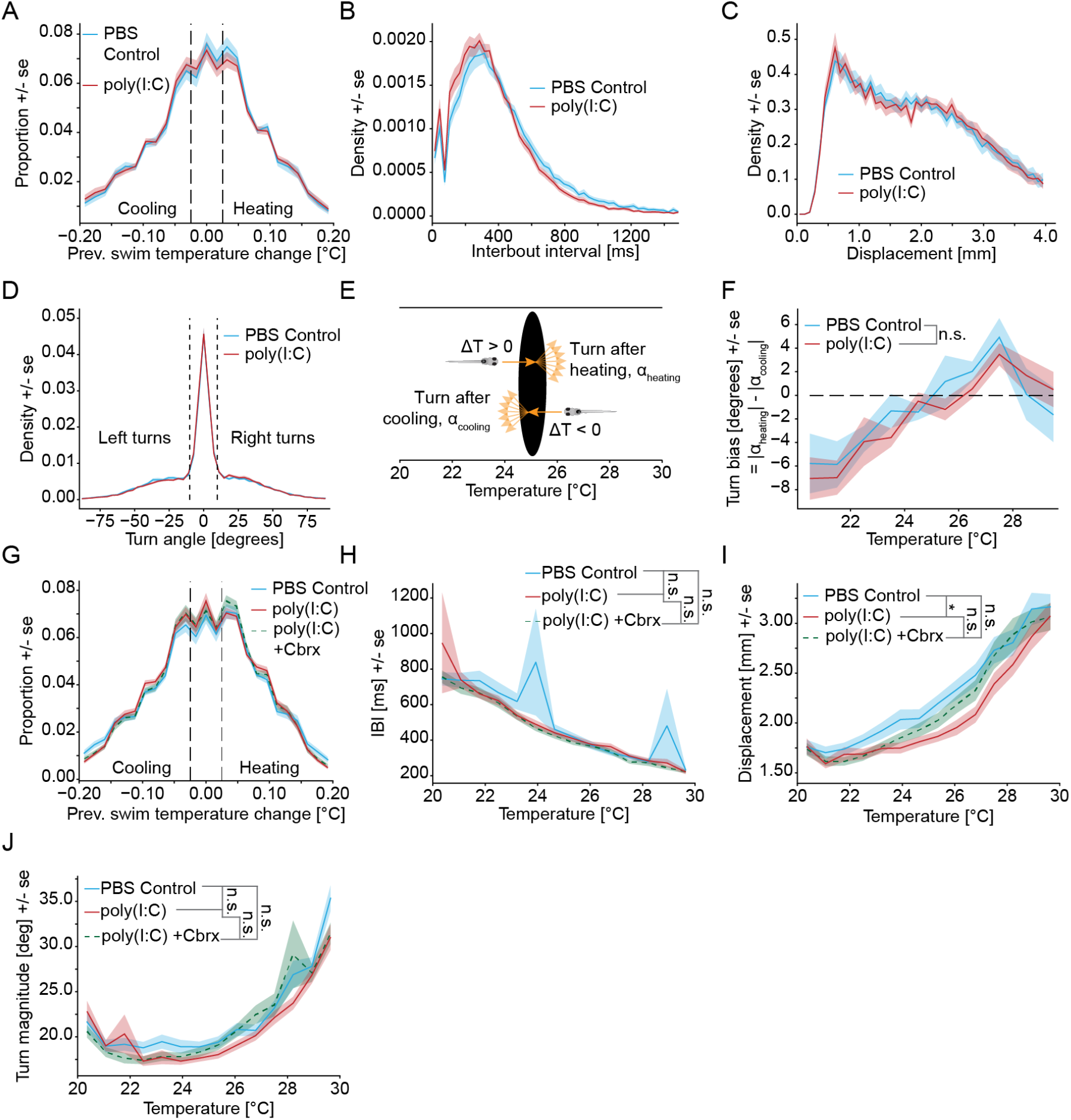
Related to Figure 4. **A**: Distribution of temperature changes experienced during individual swim bouts for PBS Control larvae (blue) and poly(I:C) larvae (red). Dashed vertical lines indicate threshold for “Heating” and “Cooling” bouts. Shading indicates bootstrap standard error across larvae. **B**: Distribution of interbout interval lengths for PBS injected controls (blue) and poly(I:C) injected larvae (red). Note that the differences in the distributions are likely a consequence of the poly(I:C) fish spending more time at warmer temperatures where interbout intervals are shorter (see Figure 1F). **C**: Distribution of swim displacements for PBS injected controls (blue) and poly(I:C) injected larvae (red). **D**: Distribution of turn angles for PBS injected controls (blue) and poly(I:C) injected larvae (red). Black dashed lines indicate threshold of 10 ° that were used to separate straight swims from turns. **E**: Illustration of the calculation of “Turn bias’. For each temperature bin, the average turn magnitude after fish have experienced heating stimuli (swimming towards warmer temperatures) and the avearge turn magnitude after fish have experienced cooling stimuli (swimming towards cooler temperatures) is calculated. Turn bias is the average magnitude after heating minus the average magnitude after cooling stimuli. I.e. positive values indicate that turns are on average larger after heating stimuli than after cooling stimuli. **F**: Turn bias at different temperatures within the gradient. Data from PBS injected control larvae in blue, from poly(I:C) injected larvae in red. Shading indicates bootstrap standard error across larvae. **G**: Distribution of temperature changes experienced during individual swim bouts for PBS Control larvae in DMSO (blue), poly(I:C) larvae in DMSO (red), and poly(I:C) larvae in Celebrex (dashed green). Dashed vertical lines indicate threshold for “Heating” and “Cooling” bouts. Shading indicates bootstrap standard error across larvae. **H**: Average interbout interval binned by gradient temperature for PBS-injected DMSO controls (blue), poly(I:C)-injected fish in DMSO (red), and poly(I:C) injected with with Celebrex (green dashed). Shading indicates bootstrap standard error across larvae. Bootstrap test of average difference across larvae. **I**: Average bout displacement binned by gradient temperature for PBS-injected DMSO controls (blue), poly(I:C)-injected fish in DMSO (red), and poly(I:C) injected with with Celebrex (green dashed). Shading indicates bootstrap standard error across larvae. Bootstrap test of average difference across larvae. **J**: Average turn magnitude binned by gradient temperature for PBS-injected DMSO controls (blue), poly(I:C)-injected fish in DMSO (red), and poly(I:C) injected with with Celebrex (green dashed). Shading indicates bootstrap standard error across larvae. Bootstrap test of average difference across larvae. In all panels, significance after Benjamini-Hochberg correction is indicated as follows. ***: p-value smaller than critical value at 0.1% FDR; **: p-value smaller than critical value at 1% FDR; *: p-value smaller than critical value at 5% FDR; n.s: p-value larger than critical value at 5% FDR. Written out p-values are uncorrected and did not pass criterion after correction.

**Figure S5:**
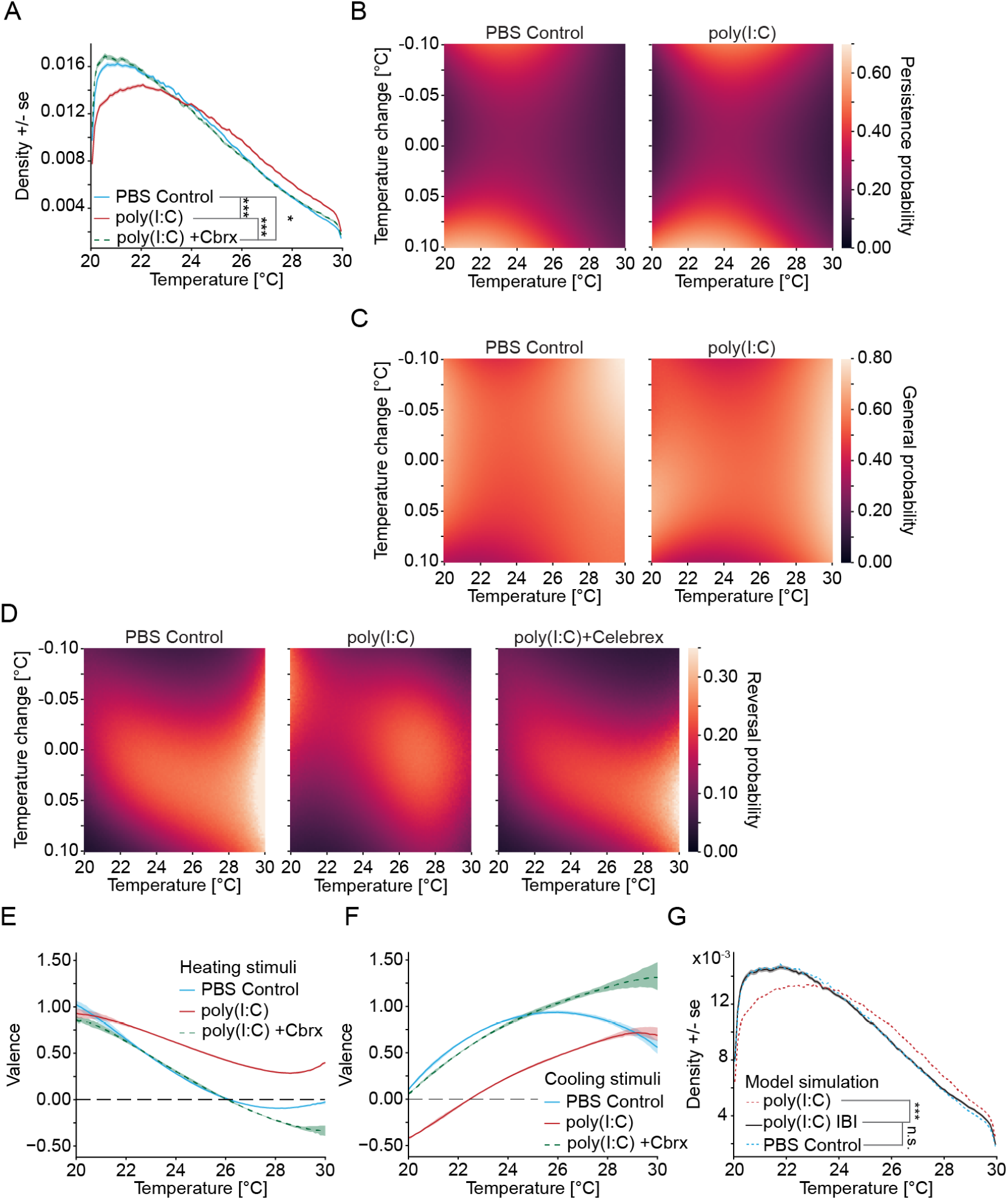
Related to Figure 5. **A**: Densities of time spent at different temperatures in gradient navigation simulations of models fit on PBS control data in DMSO (blue), poly(I:C) data in DMSO (red), and poly(I:C) data in Celebrex (green dashed). Shading indicates bootstrap standard error across simulations. **B**: Probability of being in the peristent swim mode for the two treatment conditions dependent on the current temperature (X-axis) and the temperature change across the previous bout (Y-axis). The probability is indicated with the color scale depicted to the right. For reference: Bright colors that are restricted to either cooling or heating stimuli indicate reversals that depend on the swim direction within the gradient. **C**: Same as B but for the general swim mode. **D**: Probability of being in the reversal state for the three indicated treatment conditions dependent on the current temperature (X-axis) and the temperature change across the previous bout (Y-axis). The probability is indicated with the color scale depicted to the right. For reference: Bright colors that are restricted to either cooling or heating stimuli indicate reversals that depend on the swim direction within the gradient. **E**: Valence of heating stimuli when experienced at different temperatures for PBS control larvae in DMSO (blue), poly(I:C) larvae in DMSO (red), and poly(I:C) larvae in Celebrex. Shading indiates 90% confidence intervals of the posterior distribution. **F**: Same as E but for cooling stimuli. **G**: Simulation of a hybrid model in which only the Interbout interval (IBI) parts of the emission model have been fit on poly(I:C) data (black). Original PBS control data model blue dashed, original poly(I:C) data model red dashed. In all panels, significance after Benjamini-Hochberg correction is indicated as follows. ***: p-value smaller than critical value at 0.1% FDR; *: p-value smaller than critical value at 5% FDR; n.s.: p-value larger than critical value at 5% FDR.

**Figure S6:**
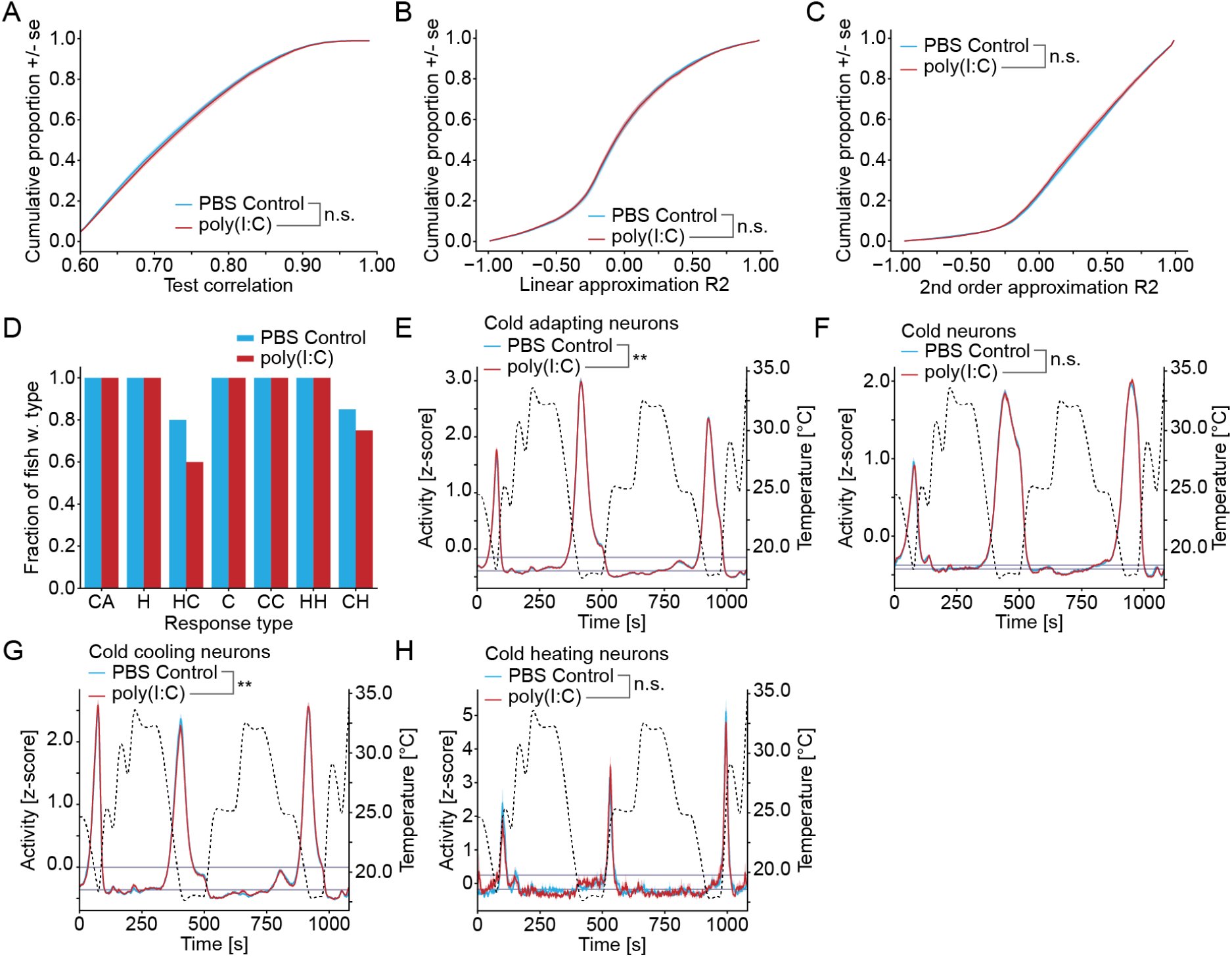
Related to Figure 6. **A**: Cumulative distribution of test-correlation values of MINE models for fit neurons. *n*.*s*. : *p >* 0.05 bootstrapped KS test across larvae. **B**: Cumulative distribution of linear approximation scores of MINE models of fit neurons. *n*.*s*. : *p >* 0.05 bootstrapped KS test across larvae. **C**: Cumulative distribution of 2nd order approximation scores of MINE models of fit neurons. *n*.*s*. : *p >* 0.05 bootstrapped KS test across larvae. **D**: For each response type the fraction of fish in which the type was identified. PBS control fish blue, poly(I:C) injected fish red. **E-H**: Comparison of cluster average responses for response types that respond in cool temperatures. Responses of PBS Control blue, Poly(I:C) red. Temperature stimulus is shown in a black dashed line (right Y-axis). Shaded error indicates bootstrap standard error of responses across larvae. Significance was determined by bootstrapping across the cluster identification procedure (see Materials and Methods). Significance after Benjamini-Hochberg correction is indicated as follows. **: p-value smaller than critical value at 1% FDR; *: p-value smaller than critical value at 5% FDR; n.s.: p-value larger than critical value at 5% FDR. Horizontal navy lines indicate computed baseline and activation thresholds used in Figure 6G. **E**: Cold adapting response type. **F**: Cold response type. **G**: Cold and cooling response type. **H**: Cold and heating response type.

## References

[1] Celsus, A. C. Cornelius celsus of medicine: In eight books (T. Chidley, London, 1837), 3 edn.

[2] Li, S., Liu, X., Lin, T. & Zhang, D. Behavioral fever in lined seahorse (hippocampus erectu) enhances the immune response to vibrio harveyi infection. Animals (Basel) 15, 1509 (2025).

[3] Sheng, Y. et al. Fruit flies exploit behavioral fever as a defense strategy against parasitic insects. Sci. Adv. 11, eadw0191 (2025).

[4] Haddad, F. et al. Fever integrates antimicrobial defences, inflammation control, and tissue repair in a cold-blooded vertebrate. Elife 12 (2023).

[5] Evans, S. S., Repasky, E. A. & Fisher, D. T. Fever and the thermal regulation of immunity: the immune system feels the heat. Nat. Rev. Immunol. 15, 335–349 (2015).

[6] Boltana, S. et al. Behavioural fever is a synergic signal amplifying the innate immune response. Proceedings of the Royal Society B: Biological Sciences 280, 20131381 (2013).

[7] Waddle, A. W. et al. Hotspot shelters stimulate frog resistance to chytridiomycosis. Nature 631, 344–349 (2024).

[8] Rakus, K., Ronsmans, M. & Vanderplasschen, A. Behavioral fever in ectothermic vertebrates. Dev. Comp. Immunol. 66, 84–91 (2017).

[9] Nakamura, K. Central circuitries for body temperature regulation and fever. Am. J. Physiol. Regul. Integr. Comp. Physiol. 301, R1207–28 (2011).

[10] Hafez, E. S. E. Behavioral thermoregulation in mammals and birds. Int. J. Biometeorol. 7, 231–240 (1964).

[11] Silva, J. E. Thermogenic mechanisms and their hormonal regulation. Physiol. Rev. 86, 435–464 (2006).

[12] Crawshaw, L. I. & Stitt, J. T. Behavioural and autonomic induction of prostaglandin E-1 fever in squirrel monkeys. J. Physiol. 244, 197–206 (1975).

[13] Briese, E. Selected temperature correlates with intensity of fever in rats. Physiol. Behav. 61, 659–660 (1997).

[14] Osterhout, J. A. et al. A preoptic neuronal population controls fever and appetite during sickness. Nature 1–8 (2022).

[15] Myhre, K., Cabanac, M. & Myhre, G. Fever and behavioural temperature regulation in the frog rana esculenta. Acta Physiol. Scand. 101, 219–229 (1977).

[16] Bernheim, H. A. & Kluger, M. J. Fever: effect of drug-induced antipyresis on survival. Science 193, 237–239 (1976).

[17] Bicego, K. C., Steiner, A. A., Antunes-Rodrigues, J. & Branco, L. G. S. Indomethacin impairs LPS-induced behavioral fever in toads. J. Appl. Physiol. 93, 512–516 (2002).

[18] Engblom, D. et al. Microsomal prostaglandin E synthase-1 is the central switch during immune-induced pyresis. Nat. Neurosci. 6, 1137–1138 (2003).

[19] Engström, L. et al. Lipopolysaccharide-induced fever depends on prostaglandin E2 production specifically in brain endothelial cells. Endocrinology 153, 4849–4861 (2012).

[20] Eskilsson, A. et al. Immune-induced fever is dependent on local but not generalized prostaglandin E2 synthesis in the brain. J. Neurosci. 37, 5035–5044 (2017).

[21] Gray, D. A., Marais, M. & Maloney, S. K. A review of the physiology of fever in birds. J. Comp. Physiol. B 183, 297–312 (2013).

[22] Quan, N. & Banks, W. A. Brain-immune communication pathways. Brain Behav. Immun. 21, 727–735 (2007).

[23] Scammell, T. E., Elmquist, J. K., Griffin, J. D. & Saper, C. B. Ventromedial preoptic prostaglandin E2 activates fever-producing autonomic pathways. J. Neurosci. 16, 6246–6254 (1996).

[24] Ilanges, A. et al. Brainstem ADCYAP1+ neurons control multiple aspects of sickness behaviour. Nature 609, 761–771 (2022).

[25] Tan, C. L. & Knight, Z. A. Regulation of body temperature by the nervous system. Neuron 98, 31–48 (2018).

[26] Haesemeyer, M., Robson, D. N., Li, J. M., Schier, A. F. & Engert, F. A brain-wide circuit model of heat-evoked swimming behavior in larval zebrafish. Neuron 98, 817–831.e6 (2018).

[27] Balakrishnan, K. A. & Haesemeyer, M. Behavioral and circuit principles of temperature gradient navigation. Curr. Biol. 35, 5395–5410.e8 (2025).

[28] Palieri, V. et al. The preoptic area and dorsal habenula jointly support homeostatic navigation in larval zebrafish. Curr. Biol. 34, 489–504.e7 (2024).

[29] Urison, N. T., Goelst, K. & Buffenstein, R. A positive fever response by a poikilothermic mammal, the naked mole rat (heterocephalus glaber). J. Therm. Biol. 18, 245–249 (1993).

[30] Anderson, L. S., Costabile, J. D., Schwinn, S., Calderon, D. & Haesemeyer, M. Sensorimotor integration enhances temperature stimulus processing. PLoS Comput. Biol. 21, e1013134 (2025).

[31] Mani, A., Haddad, F., Barreda, D. R. & Salinas, I. The telencephalon is a neuronal substrate for systemic inflammatory responses in teleosts via polyamine metabolism. Proceedings of the National Academy of Sciences 121, e2404781121 (2024).

[32] Boltana, S. et al. The expression of TRPV channels, prostaglandin E2 and pro-inflammatory cytokines during behavioural fever in fish. Brain Behav. Immun. 71, 169–181 (2018).

[33] Rey, S., Moiche, V., Boltaña, S., Teles, M. & MacKenzie, S. Behavioural fever in zebrafish larvae. Dev. Comp. Immunol. 67, 287–292 (2017).

[34] Palha, N. et al. Real-time whole-body visualization of chikungunya virus infection and host interferon response in zebrafish. PLoS Pathog. 9, e1003619 (2013).

[35] Balla, K. M., Rice, M. C., Gagnon, J. A. & Elde, N. C. Linking virus discovery to immune responses visualized during zebrafish infections. Curr. Biol. (2020).

[36] Konsman, J. P., Parnet, P. & Dantzer, R. Cytokine-induced sickness behaviour: mechanisms and implications. Trends Neurosci. 25, 154–159 (2002).

[37] Kozak, W., Conn, C. A. & Kluger, M. J. Lipopolysaccharide induces fever and depresses locomotor activity in unrestrained mice. Am. J. Physiol. 266, R125–35 (1994).

[38] Hetem, R. S. et al. Fever and sickness behavior during an opportunistic infection in a free-living antelope, the greater kudu (tragelaphus strepsiceros). Am. J. Physiol. Regul. Integr. Comp. Physiol. 294, R246–54 (2008).

[39] Shafer, M. E. R., Sawh, A. N. & Schier, A. F. Gene family evolution underlies cell-type diversification in the hypothalamus of teleosts. Nat Ecol Evol 6, 63–76 (2022).

[40] Cooper, K. E., Cranston, W. I. & Honour, A. J. Observations on the site & mode of action of pyrogens in the rabbit brain. J. Physiol. 191, 325–337 (1967).

[41] Rummel, C., Sachot, C., Poole, S. & Luheshi, G. N. Circulating interleukin-6 induces fever through a STAT3-linked activation of COX-2 in the brain. Am. J. Physiol. Regul. Integr. Comp. Physiol. 291, R1316–26 (2006).

[42] Eskilsson, A., Tachikawa, M.Hosoya, K.-I. & Blomqvist, A. Distribution of microsomal prostaglandin E synthase-1 in the mouse brain. J. Comp. Neurol. 522, 3229–3244 (2014).

[43] Breder, C. D. & Saper, C. B. Expression of inducible cyclooxygenase mRNA in the mouse brain after systemic administration of bacterial lipopolysaccharide. Brain Res. 713, 64–69 (1996).

[44] Eskilsson, A. et al. Immune-induced fever is mediated by IL-6 receptors on brain endothelial cells coupled to STAT3dependent induction of brain endothelial prostaglandin synthesis. J. Neurosci. 34, 15957–15961 (2014).

[45] Nadeau, S. & Rivest, S. Effects of circulating tumor necrosis factor on the neuronal activity and expression of the genes encoding the tumor necrosis factor receptors (p55 and p75) in the rat brain: a view from the blood-brain barrier. Neuroscience 93, 1449–1464 (1999).

[46] Ushikubi, F. et al. Impaired febrile response in mice lacking the prostaglandin E receptor subtype EP3. Nature 395, 281–284 (1998).

[47] Lazarus, M. et al. EP3 prostaglandin receptors in the median preoptic nucleus are critical for fever responses. Nat. Neurosci. 10, 1131–1133 (2007).

[48] Machado, N. L. S., Bandaru, S. S., Abbott, S. B. G. & Saper, C. B. EP3R-expressing glutamatergic preoptic neurons mediate inflammatory fever. J. Neurosci. 40, 2573–2588 (2020).

[49] Yahiro, T., Nakamura, Y. & Nakamura, K. The pyrogenic mediator prostaglandin E(2) elicits warmth seeking via EP3 receptor-expressing parabrachial neurons: a potential mechanism of chills. J. Physiol. (2026).

[50] Sugimoto, Y. & Narumiya, S. Prostaglandin E receptors *. J. Biol. Chem. 282, 11613–11617 (2007).

[51] Vasilache, A. M., Andersson, J. & Nilsberth, C. Expression of PGE2 EP3 receptor subtypes in the mouse preoptic region. Neurosci. Lett. 423, 179–183 (2007).

[52] Wang, X. et al. The lateral habenula contributes to regulation of body temperature. iScience 28, 112923 (2025).

[53] Gong, L. et al. Celecoxib pathways: pharmacokinetics and pharmacodynamics. Pharmacogenet. Genomics 22, 310–318 (2012).

[54] McNab, F., Mayer-Barber, K., Sher, A., Wack, A. & O’Garra, A. Type I interferons in infectious disease. Nat. Rev. Immunol. 15, 87–103 (2015).

[55] Altan, E. et al. A highly divergent picornavirus infecting the gut epithelia of zebrafish (danio rerio) in research institutions worldwide. Zebrafish 16, 291–299 (2019).

[56] Le Goc, G. et al. Thermal modulation of zebrafish exploratory statistics reveals constraints on individual behavioral variability. BMC Biol. 19, 208 (2021).

[57] Herrera, K. J., Panier, T., Guggiana-Nilo, D. & Engert, F. Larval zebrafish use olfactory detection of sodium and chloride to avoid salt water. Curr. Biol. 0 (2020).

[58] Geyer, C. J. Practical markov chain monte carlo. Stat. Sci. 7, 473–483 (1992).

[59] Paoli, E., Palieri, V., Shenoy, A. & Portugues, R. Modulation of habenula axon terminals supports action-outcome associations in larval zebrafish. bioRxiv 2025.02.13.638047 (2025).

[60] Pachitariu, M. et al. Suite2p: beyond 10,000 neurons with standard two-photon microscopy. bioRxiv 061507 (2017).

[61] Costabile, J. D., Balakrishnan, K. A., Schwinn, S. & Haesemeyer, M. Model discovery to link neural activity to behavioral tasks. Elife 12 (2023).

[62] Bronstein, S. M. & Conner, W. E. Endotoxin-induced behavioural fever in the madagascar cockroach, gromphadorhina portentosa. J. Insect Physiol. 30, 327–330 (1984).

[63] Casterlin, M. E. & Reynolds, W. W. Behavioral fever in anuran amphibian larvae. Life Sci. 20, 593–596 (1977).

[64] Akins, C., Thiessen, D. & Cocke, R. Lipopolysaccharide increases ambient temperature preference in C57BL/6J adult mice. Physiol. Behav. 50, 461–463 (1991).

[65] Ranels, H. J. & Griffin, J. D. The effects of prostaglandin E2 on the firing rate activity of thermosensitive and temperature insensitive neurons in the ventromedial preoptic area of the rat hypothalamus. Brain Res. 964, 42–50 (2003).

[66] Cutler, B. & Haesemeyer, M. Vertebrate behavioral thermoregulation: knowledge and future directions. Neurophotonics 11, 033409 (2024).

[67] Nakamura, K. et al. The rostral raphe pallidus nucleus mediates pyrogenic transmission from the preoptic area. J. Neurosci. 22, 4600–4610 (2002).

[68] Nagashima, K., Nakai, S., Tanaka, M. & Kanosue, K. Neuronal circuitries involved in thermoregulation. Auton. Neurosci. 85, 18–25 (2000).

[69] Bicego, K. C. & Branco, L. G. S. Discrete electrolytic lesion of the preoptic area prevents LPS-induced behavioral fever in toads. J. Exp. Biol. 205, 3513–3518 (2002).

[70] Liu, R. et al. Lateral habenula neurons signal cold aversion and participate in cold aversion. Neurochem. Res. (2023).

[71] Hikosaka, O. The habenula: from stress evasion to value-based decision-making. Nat. Rev. Neurosci. 11, 503–513 (2010).

[72] Sylwestrak, E. L. et al. Cell-type-specific population dynamics of diverse reward computations. Cell 185, 3568–3587.e27 (2022).

[73] Turner, K. J. et al. Afferent connectivity of the zebrafish habenulae. Front. Neural Circuits 10, 30 (2016).

[74] Garland, J. C. & Mogenson, G. J. An electrophysiological study of convergence of entopeduncular and lateral preoptic inputs on lateral habenular neurons projecting to the midbrain. Brain Res. 263, 33–41 (1983).

[75] Kowski, A. B., Geisler, S., Krauss, M. & Veh, R. W. Differential projections from subfields in the lateral preoptic area to the lateral habenular complex of the rat. J. Comp. Neurol. 507, 1465–1478 (2008).

[76] Westerfield, M. The zebrafish book. A guide for the laboratory use of zebrafish (Danio rerio) (Univ. of Oregon Press, Eugene, Oregon, 2000), 4 edn.

[77] Shainer, I. et al. A single-cell resolution gene expression atlas of the larval zebrafish brain. Sci. Adv. 9, eade9909 (2023).

[78] Lovett-Barron, M. et al. Multiple convergent hypothalamus-brainstem circuits drive defensive behavior. Nat. Neurosci. 23, 959–967 (2020).

[79] Rohlfing, T. & Maurer, C. R., Jr. Nonrigid image registration in shared-memory multiprocessor environments with application to brains, breasts, and bees. IEEE Trans. Inf. Technol. Biomed. 7, 16–25 (2003).

[80] Carpenter, B. et al. Stan: A probabilistic programming language. J. Stat. Softw. 76, 1–32 (2017).

[81] Miri, A., Daie, K., Burdine, R. D., Aksay, E. & Tank, D. W. Regression-based identification of behavior-encoding neurons during large-scale optical imaging of neural activity at cellular resolution. J. Neurophysiol. 105, 964–980 (2011).

[82] Hao, Y. et al. Integrated analysis of multimodal single-cell data. Cell 184, 3573–3587.e29 (2021).

